# ATRX Deficiency Drives Aberrant Type I Interferon Signalling Through cGAS-Dependent Transcriptional Dysregulation

**DOI:** 10.1101/2025.11.19.685761

**Authors:** Marie-Thérèse El-Daher, Eve L White, Graeme Grimes, Gaofeng Zhu, Andreas Fellas, Tom A Tait, Carolina Uggenti, Natalie Blair, Erisa Nita, Philippe Gautier, Hywel Dunn-Davies, Sophie Glen, Somdutta Dhir, Mathieu P Rodero, Gillian I Rice, Luis Seabra, Richard Clark, Angie Fawkes, Joseph A Marsh, John H Livingston, Evangeline Wassmer, Swati Naik, Julie Vogt, Bertrand Isidor, Rebekah Tillotson, Alice Lepelley, Patrick Revy, Richard J Gibbons, Yanick J Crow

## Abstract

The X-linked α-thalassaemia intellectual disability syndrome (ATRX) protein is a chromatin remodeller involved in transcriptional regulation and genome stability. While the importance of ATRX in development and malignancy is well recognised, its role in innate immunity is less well defined. In two unrelated patients demonstrating cerebral white matter disease, learning difficulties and a persistent upregulation of interferon stimulated gene expression in whole blood, we identified the same Y1758C missense substitution in ATRX. Using patient-derived cells, engineered fibroblasts and neuronal models, we show that this substitution, and other loss of function mutations in ATRX, result in enhanced type I interferon signalling through a cGAS-dependent mechanism uncoupled from the DNA sensing activity of cGAS. Loss of ATRX function leads to alterations in the chromatin distribution of DAXX and H3.3, with cGAS essential for the changes in nucleosome composition and gene expression mediated by ATRX deficiency. Thus, our study highlights a previously unrecognized link between ATRX dysfunction and inflammation involving a non-canonical role of cGAS.

## INTRODUCTION

The X-linked α-thalassaemia intellectual disability syndrome (ATRX) protein acts as an ATP-dependent molecular motor involved in chromatin remodelling and transcriptional regulation^1^. Positioned predominantly in the nucleus, ATRX exhibits a broad genomic distribution, localizing to heterochromatic features including silenced repetitive elements, telomeres and pericentromeres, and to euchromatin regions^2^. In complex with death domain associated protein (DAXX), ATRX facilitates the deposition of the histone variant H3.3 at sites of nucleosome turnover in a replication-independent manner^3^, and as a key epigenetic regulator, disturbance of ATRX function can impact DNA replication, chromatin state and gene expression^4^. Germline mutations in *ATRX* cause a sex-linked recessive disorder associated with learning difficulties, referred to as ATR-X syndrome^5, 6^. Furthermore, acquired somatic loss of function mutations in ATRX are characteristic of tumours that utilize the alternative lengthening of telomeres (ALT) pathway, an adaptive mechanism active in 10-15% of all cancers^7, 8^. Notably, mutations in ATRX/DAXX complex are present in 90% of ALT-positive gliomas, frequently co-occurring with mutations in isocitrate dehydrogenase (IDH)^9^, the latter facilitating tumour immune evasion by limiting the inflammatory response triggered by ATRX deficiency^10^. ATRX also contributes to innate immune defence by restricting viral gene expression through the promotion of heterochromatin formation^11^. These multiple functions of ATRX underscore its importance not only in development and cancer biology, but also in innate immunity.

While type I interferons are integral to the response to viral infection, establishing an anti-viral state through the upregulation of hundreds of interferon stimulated genes (ISGs)^12^, inappropriate activation of an interferon response can be pathogenic, leading to tissue damage. This dichotomy is particularly well illustrated by the type I interferonopathies, Mendelian diseases characterised by chronically enhanced type I interferon signalling^13^. Type I interferons can be induced through the sensing of both foreign and self-nucleic acids by innate immune receptors, among them cyclic GMP-AMP synthase (cGAS)^14^. Upon binding to double-stranded DNA in a sequence-independent manner, cGAS catalyses the production of the second messenger 2′-3′ cyclic guanosine monophosphate-adenosine monophosphate (cGAMP) from GTP and ATP. cGAMP binds to the signalling adaptor stimulator of interferon genes (STING). Activated STING is transported from the endoplasmic reticulum (ER) to the ER-Golgi intermediate compartment and the Golgi where it recruits Tank binding kinase 1 (TBK1), which in turn activates interferon regulatory factor 3 (IRF3) and nuclear factor kappa B subunit (NFκB). Both of these transcription factors then translocate to the nucleus, leading to the induction of type I interferon and other pro-inflammatory cytokine genes^15^. Sensing of self-DNA by cGAS has been shown to occur in a number of type I interferonopathies, for example due to loss-of-function mutations in the three prime repair exonuclease TREX1^16^. While originally considered as an exclusively cytoplasmic protein, it is now recognised that cGAS also localises to the nucleus, where it is associated with nucleosomes, DNA replication forks, double-strand breaks and centromeres^17, 18^. These observations suggest that cGAS may have functions beyond its canonical role in cytosolic DNA sensing. In this study, we report that loss of ATRX induces type I interferon signalling and neuroinflammation, a process that depends on the interplay between the ATRX/DAXX complex, H3.3 and cGAS in regulating transcription.

## RESULTS

### A Y1758C substitution in ATRX causes neuroinflammation consistent with a type I interferonopathy

As part of an ongoing protocol involving the agnostic screening of patients with uncharacterized disease phenotypes for an upregulation of type I interferon signalling, we identified two unrelated patients (P1 and P2, from families one and two, respectively) (Fig. 1A) demonstrating similar cerebral white matter lesions and intracranial calcification, moderate to severe intellectual disability, and facial features consistent with ATR-X syndrome in P2 (Fig. 1B, fig. S1A-C). Exome sequencing in P1 and P2 revealed the same p.(Tyr1758Cys) (Y1758C, c.5273A>G) substitution in the helicase domain of ATRX, which was inherited from one parent in the absence of a family history of ATR-X syndrome or inflammatory disease in both cases. Two relatives of P1 were found not to carry the variant (Fig. 1A). This substitution is not present on gnomAD v4 (0/1,018,707 alleles) and is predicted as pathogenic by *in silico* analyses, with the tyrosine residue at position 1758 conserved to C. elegans (Fig. 1C). Clinically accredited genome-wide DNA methylation testing in P2 revealed an episignature typical of ATR-X syndrome^19^. As evidence of enhanced interferon signalling, ISG expression in whole blood was persistently elevated in both individuals on all occasions tested (six times in P1, and eight in P2) (Fig. 1D, fig. S1D-F). Cerebral white matter disease, reminiscent of type I interferon driven neuroinflammation^20^, has been reported in other patients with ATR-X syndrome^21, 22^. By mining publicly available RNA sequencing (RNA-seq) datasets relating to ATRX^23, 24, 25^, and performing gene set enrichment pathway analysis, we identified evidence for an upregulation of interferon responses in *ATRX* null cells (figure S2). These observations prompted us to further explore interferon signalling in the context of ATRX-related disease.

**Figure 1.**
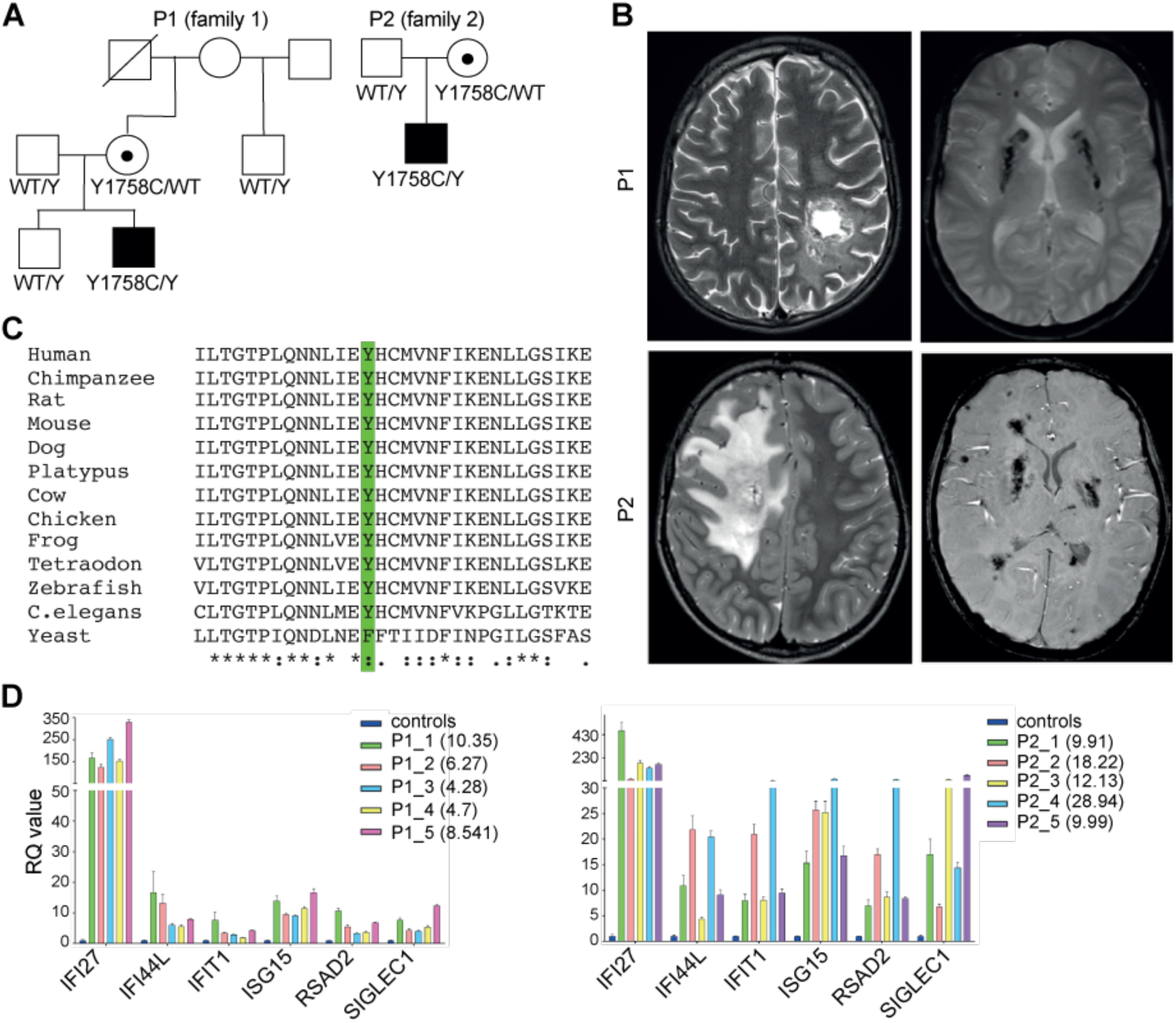
ATRX-related type I interferonopathy. (A) Pedigrees of P1 and P2 families with the X-linked variant c.5273A>G, p.Y1758C in ATRX; WT = wild-type. (B) Representative cerebral MRI images demonstrating tumefactive lesions (top and bottom left: axial T2 weighted) and dense calcification (right: axial SWI in P1 and GRE in P2). (C) CLUSTAL Omega alignment of ATRX homologs. The position of the substituted Y1758 is indicated in green. (D) Relative quantification (RQ) values of six interferon stimulated genes (ISGs) measured on five different occasions in whole blood of P1 (left) and P2 (right). Number in brackets is the interferon score calculated from the median fold change in RQ value for the panel of six ISGs indicated on the x axis (normal <; 2.4). Colours relate to different dates at sampling. Dark blue bars represent the composite data of 29 controls.

### Loss of function of ATRX results in enhanced type I interferon signalling

Germline loss of function mutations in *ATRX* are non-viable^26^, so that mutations causing ATR-X syndrome are considered to act as hypomorphs. All previously analysed mutations result in reduced ATRX protein levels to 7-50% of controls^27, 28, 29^, with some also demonstrated to impact binding to heterochromatin or chromatin remodelling activity^28, 29^. Consistent with these observations, levels of ATRX protein were reduced in primary fibroblasts obtained from four patients (P3 - P6). In contrast, ATRX levels were normal in primary fibroblasts from P1 and P2 (Fig. 2A). In primary fibroblasts from both P1 and P2, we recorded interferon score, based on the expression of a panel of 7 representative ISGs, and the expression of interferon beta gene (*IFNB1*) to be increased (Fig. 2B, 2C). The interferon score and *IFNB1* expression were also increased in primary fibroblasts from P5 and P6, but not P3 and P4 (Fig. 2B-C). Given the potential variability of genetic background in available patient material, we introduced the Y1758C mutation (referred to as knock-in (KI)) into BJ-5ta immortalised fibroblasts (referred to as BJ-hTERT), and also generated *ATRX* knock-out (KO) clones using CRISPR/Cas9 technology. As in patient primary fibroblasts, ATRX protein levels were unaffected in BJ-hTERT KI clones, while we could not detect ATRX protein in the engineered KO clones (Fig. 2D). ISG and *IFNB1* expression was elevated in both KI and KO *ATRX* clones (Fig. 2E-F). Transcriptomic analysis by RNA-seq of three independent KI, KO and WT clones revealed 4529 and 3035 differentially expressed genes (adjusted p value < 0.1) in *ATRX* KI and KO cells, respectively compared to WT clones (Fig. 2G). 2075 genes were similarly dysregulated in both KI and KO *ATRX* cells, suggesting that, at least in part, the Y1758C mutation recapitulates ATRX loss of function. Gene set enrichment analysis of these expression data identified significant upregulation of the interferon response, and of genes involved in epithelial mesenchymal transition (Fig. 2H). Concordant with these data, short-hairpin (sh) RNA mediated knock-down of *ATRX* in the human monocytic cell line THP-1 resulted in increased ISG expression and type I interferon production (fig. S3A-C).

**Figure 2.**
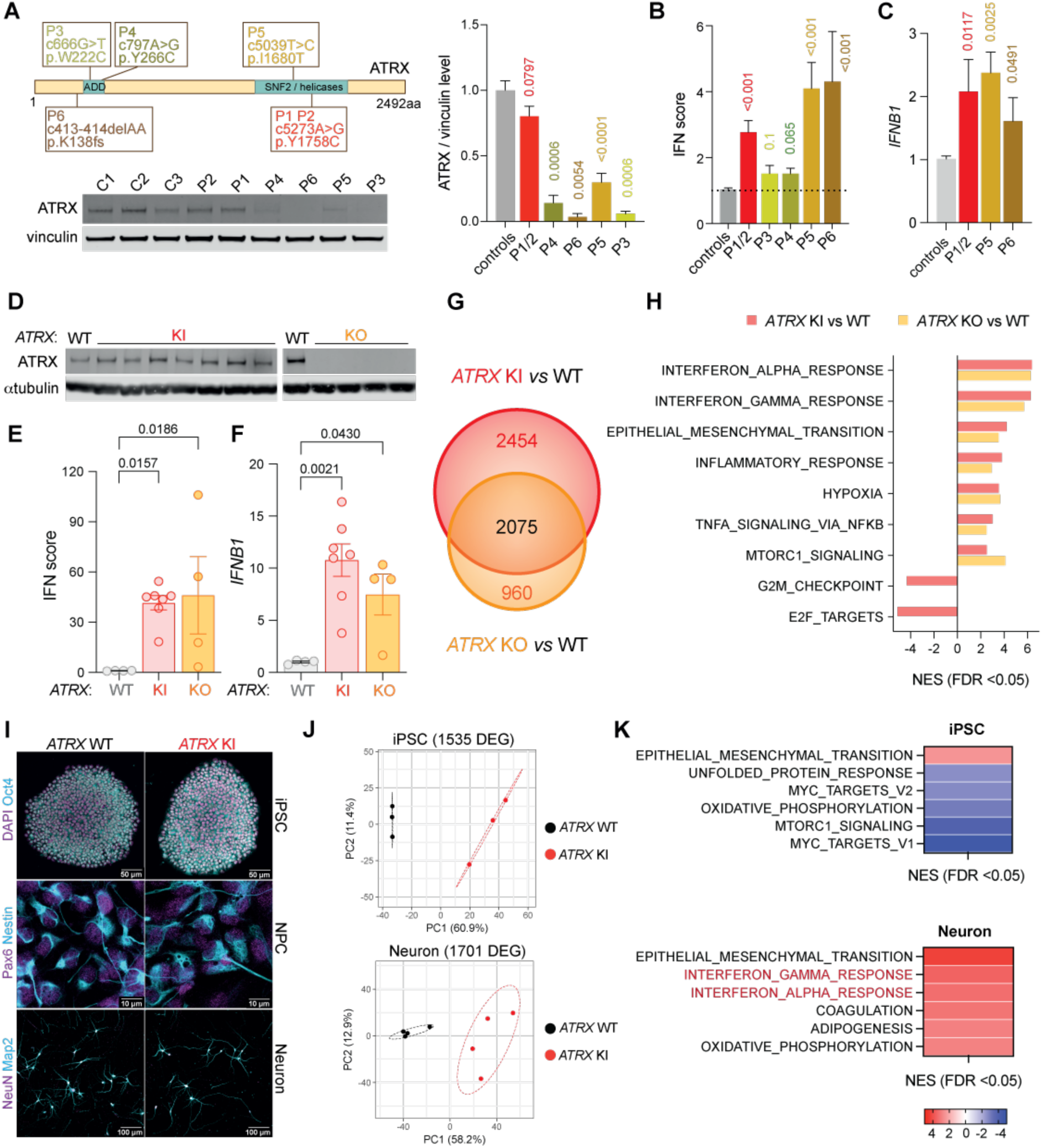
Enhanced interferon signalling in ATRX mutated cells. (A) Schematic of the ATRX protein (top left) illustrating the Y1758C mutation in P1 and P2, and the mutations in P3 – P6; ADD = ATRX-DNMT3-DNMTL-like domain (amino acid (aa) residues 161-292; SNF2 (Sucrose Non-Fermentable) domain encompassing ATPase/Helicase domains, residues 1550-2226. Representative western blot (bottom left) and quantification (right) of ATRX protein in primary fibroblasts from controls (C1 – C3) and P1 – P6, [mean of three independent experiments for P3, and 4 independent experiments for the other patients; levels were compared to controls using a Kruskal-Wallis test, two-stage linear step-up procedure of Benjamin, Krieger and Yekutieli with resulting q values given]. (B) Interferon (IFN) score calculated by qPCR representing the median of the relative expression levels of IFI27, RSAD2, MX1, ISG15, IFI44L, IFIT1, TNFAIP3 in primary fibroblasts from ATR-X syndrome patients and three controls, [six to 10 independent experiments for each sample; mean +/-SEM; scores were compared to controls using a Kruskal-Wallis test, uncorrected Dunn’s test, and resulting q values given.]. (C) Relative expression level of IFNB1 in primary fibroblasts [three to four independent experiments, levels were compared to controls using a Mann-Whitney test with resulting q values given]. (D) ATRX protein expression assessed by western blot in BJ-hTERT clones knock-in (KI) for the Y1758C mutation or knock-out (KO) for ATRX. (E) IFN score (median of the relative expression of the following ISGs: IFI27, RSAD2, MX1, ISG15, IFI44L, IFIT1, TNFAIP3) in BJ-hTERT clones WT, KI or KO for ATRX measured by qPCR. (F) Relative expression of IFNB1 in clones as in (E). [E, F: each circle represents the mean of a clone, clones were tested two to four times; Kruskal-Wallis test, two-stage linear step-up procedure of Benjamin, Krieger and Yekutieli]. (G) Differential gene expression analysis comparing three independent BJ-hTERT clones WT, KI or KO for ATRX. Top and bottom values represent the number of genes exclusively dysregulated in cells KI or KO for ATRX, respectively. Middle value represents the number of genes dysregulated in both KI and KO ATRX cells compared to WT. (H) Bar plots depicting normalised enrichment scores of hallmark gene sets with a False Discovery Rate (FDR) <; 0.05, comparing differentially expressed genes identified between WT and KI or WT and KO ATRX cells. (I) Immunofluorescence of induced pluripotent stem cells (iPSCs), iPSC-derived neural progenitor cells (NPCs) and forebrain neurons in isogenic WT or KI ATRX mutant cells. (J) Principal component analysis of RNA-seq data derived from iPSCs and forebrain neurons. DEG refers to genes differentially expressed between WT and KI ATRX cells. (K) Heatmaps depicting normalised enrichment scores (NES) of hallmark gene sets with FDR <; 0.05.

Since ATR-X syndrome is primarily a disorder of the brain, we next introduced the Y1758C mutation into induced pluripotent stem cells (iPSCs). iPSCs KO for *ATRX* did not survive clonal expansion and hence could not be studied. Using three WT and three KI clones, we then differentiated iPSCs to neural progenitor cells (NPCs) and forebrain neurons (fig. S3D). Immunostaining and global mRNA expression analysis using 3’ end RNA-seq confirmed the expression of expected iPSC, NPC and neuronal lineage markers (Fig. 2I, fig. S3E). Mutation of *ATRX* led to marked differences in gene expression in both iPSCs and neurons (Fig. 2J), with functional pathway enrichment analysis identifying significant dysregulation of several transcriptional programmes, including the emergence of an interferon response in Y1758C KI neurons (Fig. 2K).

These observations indicate that, as in patient blood and patient-derived primary fibroblasts, multiple cell types mutated or null for *ATRX* demonstrate activation of type I interferon signalling.

### cGAS is required for the activation of interferon signalling in ATRX mutant cells

The induction of type I interferon signalling involves host-encoded nucleic acid-binding pattern-recognition receptors. These include the cytosolic sensors of dsRNA (melanoma differentiation associated protein 5, MDA5, and retinoic acid inducible gene I, RIG-I), which signal through mitochondrial antiviral signalling protein (MAVS)^30^, and dsDNA sensors, most particularly cGAS, which signals to STING through the second messenger cGAMP^31^. Both RNA and DNA sensing pathways activate IRF3, leading to the subsequent induction of type I interferons expression. In turn, type I interferons are secreted and bind to the interferon-α/β receptor (IFNAR), initiating a JAK-STAT signalling cascade and phosphorylation process that induces the expression of ISGs. We detected increased levels of phosphorylated STAT1, a marker of interferon signalling through IFNAR, in both *ATRX* KI and KO cells, in the absence of any additional stimulation, and in control cells stimulated with the STING agonist diABZI (Fig. 3A). Exposing *ATRX* mutant cells to anifrolumab, a monoclonal antibody that blocks IFNAR signalling, led to the reduced expression of 35 of the 183 ISGs dysregulated in these cells prior to treatment (Fig. 3B), indicating a contribution of IFNAR downstream signalling in ATRX-mediated inflammation.

**Figure 3.**
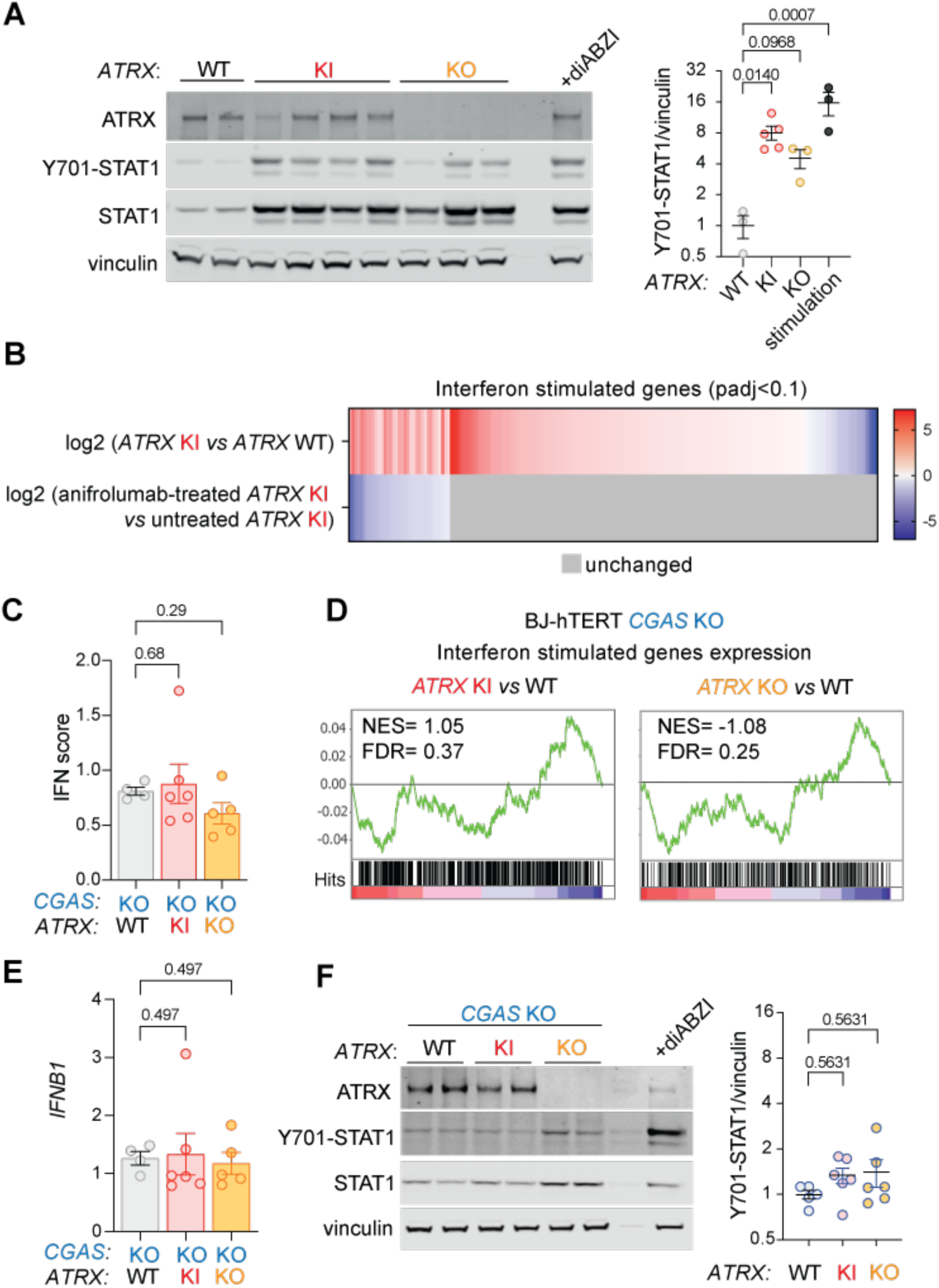
cGAS is required for the activation of interferon signalling in ATRX deficient cells. (A) Left: immunoblot of ATRX, phosphorylated (Y701) and total STAT1, and vinculin in individual BJ-hTERT fibroblast clones WT, KI (Y1758C) or KO for *ATRX*. The last lane relates to *ATRX* WT cells treated with the STING agonist diABZI. Right: quantification of phosphorylated STAT1 as in (A); ‘stimulation’ indicates cells WT for *ATRX* treated with HT-DNA or diABZI, [each dot represents the mean of a clone, except for WT in which the three dots represent two clones and the WT parental line. Clones were tested one to four times; black bars: mean +/-SEM; Unpaired t test]. (B) Effect of treatment with anifrolumab, an IFNAR inhibitor, on ISG expression in BJ-hTERT WT or KI for *ATRX*. The heatmap depicts the log2 fold change in gene expression of KI versus WT cells in untreated (top row) and treated (bottom row) cells. Gene expression was determined by RNA-seq and DESeq2 analysis. For the untreated condition, only ISGs differentially expressed between WT and KI cells are displayed (adjusted p value <; 0.1). Upon treatment, genes whose expression decreased are indicated in blue, and genes showing no change in expression in grey. (C) Interferon (IFN) score (median of the relative expression of the following ISGs: *IFI27, RSAD2, MX1, ISG15, IFI44L, IFIT1, TNFAIP3*) in BJ-hTERT KO for *CGAS* and either WT, KI, or KO for *ATRX* [C and E: each circle represents the mean of a clone, clones were tested two to four times; Kruskal-Wallis test, two-stage linear step-up procedure of Benjamin, Krieger and Yekutieli]. (D) Gene set enrichment analysis comparing the expression levels of 444 canonical ISGs determined by RNA-seq in BJ-hTERT fibroblasts lacking *CGAS*, and KI (top panel) or KO (bottom panel) for *ATRX*, relative to WT. NES = Normalized Enrichment Score; FDR = False Discovery Rate. (E) Interferon beta (*IFNB1*) gene expression as in (C). (F) Same as in (A) but in BJ-hTERT cells KO for *CGAS* [each dot represents the mean of a clone; clones were tested one to four times; black bars: mean +/-SEM; Unpaired t test].

Next, we investigated the role of discrete nucleic acid sensors in the observed upregulation of interferon signalling. ISG expression remained elevated following *MAVS* depletion by siRNA in primary fibroblasts from patients with *ATRX* mutations (fig. S4A-B) and in THP-1 cells KO for *MAVS* and knocked-down for *ATRX* (fig. S4C-D), suggesting that cytosolic RNA sensing is not involved in the interferon response induced by ATRX deficiency. To interrogate the role of DNA sensing, we examined the impact of ATRX loss of function in BJ-hTERT fibroblasts null for *CGAS*^32, 33^ (fig. S4E), and in CRISPR-engineered clones either KI or KO for *ATRX*. Notably, the increased expression of a curated list of 444 canonical ISGs observed in ATRX deficient cells was absent in cells also KO for *CGAS* (Fig. 3C-D). Of further note, while ATRX loss of function induced *IFNB1* upregulation and STAT1 phosphorylation in WT cells (Fig. 2F, 3A), *IFNB1* expression and STAT1 phosphorylation in cells KI or KO for *ATRX* were comparable to cells WT for *ATRX* on a cGAS null background (Fig. 3E-F). Consistent with these observations, genome-wide transcriptomic analysis revealed little variation between WT cells and ATRX deficient cells on a *CGAS* KO background (fig. S4F), in contrast to cells WT for *CGAS* (fig. S4G).

Collectively, these results demonstrate a requirement for cGAS in mediating the enhanced interferon response observed in the context of ATRX deficiency.

### While interferon induction in ATRX deficient cells is cGAS-dependent, this does not involve the canonical DNA sensing pathway

Having demonstrated that ATRX deficiency leads to enhanced interferon signalling dependent on cGAS, we next investigated the activation of the canonical sensing pathway downstream of cGAS, first analysing levels of its product, the second messenger cGAMP^34, 35^. Despite sample concentrations falling within the reliable detection range of the ELISA measurements (fig. S5A), cGAMP levels in primary fibroblasts from patients with ATRX mutations were in the control range (Fig. 4A). In contrast, levels of cGAMP were elevated in fibroblasts from patients with the exemplar cGAS-mediated type I interferonopathies, Aicardi-Goutières syndrome (here, due to mutations in TREX1 and RNASEH2A), and DNase II deficiency (Fig. 4A). While demonstrating the expected reactivity to DNA stimulation, cGAMP levels were also found to be in the WT range in ATRX mutant BJ-hTERT fibroblasts (Fig 4B, fig. S5B). Correspondingly, we observed no difference in the phosphorylation status of effectors involved in type I interferon induction downstream of cGAS activation by DNA and cGAMP production (and upstream of IFNAR and STAT1 phosphorylation), i.e., STING, TBK1, IRF3 and NF-κB p65, in *CGAS* WT BJ-hTERT clones mutant for ATRX compared to control cells (Fig. 4C). Additionally, because cGAS resides in the nucleus and its detachment from chromatin could trigger DNA sensing, we verified that cGAS remained tightly tethered to chromatin in ATRX deficient cells, similar to control cells (Fig. 4D).

**Figure 4.**
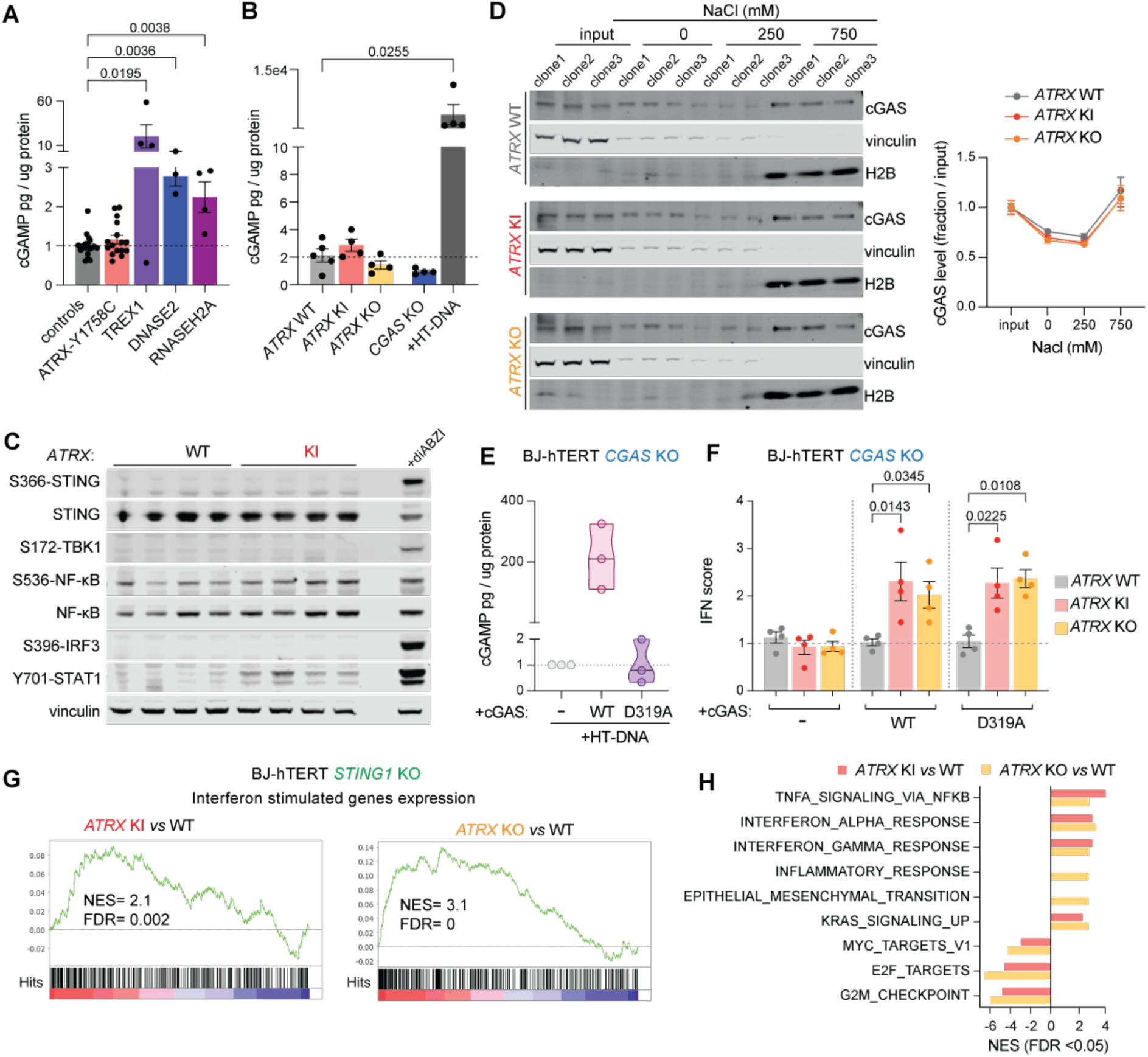
cGAS, but not its second messenger 2’3’cGAMP or STING, is required for the activation of interferon stimulated genes in ATRX deficient cells. (A) cGAMP measured by ELISA in unstimulated control fibroblasts, ATRX patient fibroblasts (Y1758C), and fibroblasts from patients with biallelic mutations in *TREX1*, *DNASE2* or *RNASEH2A* [circles represent technical repeats: 19 repeats in controls, 16 repeats in ATRX-Y1758C patients, and three or four repeats in patients with mutations in *TREX1*, *DNASE2* or *RNASEH2A*. Kruskal-Wallis test, uncorrected Dunn’s test, mean +/-SEM]. (B) cGAMP levels measured by ELISA in unstimulated BJ-hTERT fibroblasts WT for *CGAS* and either WT, KI (Y1758C) or KO for *ATRX*, and in *CGAS* KO BJ-hTERT and WT fibroblasts treated with herring-testis DNA (HT-DNA), [each circle represents the mean of a clone, clones were tested two to four times; Kruskal-Wallis test, uncorrected Dunn’s test, mean +/-SEM]. (C) Immunoblot of phosphorylated STING (S366), phosphorylated and total forms of NF-κB, phosphorylated IRF3 (S396), TBK1 (S172) and STAT1 (Y701) in individual clones of BJ-hTERT fibroblasts WT or KI for *ATRX*. The last lane relates to WT BJ-hTERT fibroblasts treated with the STING agonist diABZI, mimicking cGAMP activity. (D) BJ-hTERT fibroblasts WT, KI or KO for *ATRX* were separated into cytosolic and nuclear fractions, followed by sequential elution of nuclear pellets with the indicated concentrations of NaCl. Quantification (right) of cGAS levels in the different fractions of cells WT, KI or KO for *ATRX*. Vinculin and H2B are cytosolic and chromatin markers respectively [three independent clones, mean +/-SEM]. (E) cGAMP levels measured in BJ-hTERT fibroblasts KO for *CGAS* transfected with lipofectamine only (-), WT or catalytically inactive (D319A mutant) cGAS constructs and stimulated with HT-DNA for four hours. (F) Interferon (IFN) score (median of the relative expression of the following ISGs: *IFI27, IFI44L, OASL, IFIT2, IFI6, ALDHA1*) in BJ-hTERT fibroblasts KO for *CGAS*, and WT, KI or KO for *ATRX* were transfected with lipofectamine only (-), WT or catalytic inactive (D319A mutant) cGAS constructs, [each circle represents the mean of a clone, clones were tested four times, average+/-SEM, Kruskal-Wallis test, uncorrected Dunn’s test]. (G) Gene set enrichment analysis (GSEA) comparing the expression levels of 444 canonical ISGs determined by RNA-seq in BJ-hTERT fibroblasts lacking *STING1*, and KI (top panel) or KO (bottom panel) for *ATRX*, relative to WT. NES = Normalized Enrichment Score; FDR = False Discovery Rate. (H) Bar plots depicting NES of hallmark gene sets that were significantly different (FDR <; 0.05) in BJ-hTERT *STING1* KO fibroblasts, comparing cells KI or KO for *ATRX* to cells WT for *ATRX*.

The finding that cGAS is present in the nucleus has raised the possibility of additional roles for cGAS distinct from its DNA sensing function^17, 18^. To further establish the non-requirement for cGAS-dependent cGAMP production in mediating the ATRX inflammatory phenotype, we overexpressed WT or catalytically dead (D319A mutant) cGAS in BJ-hTERT fibroblasts null for *CGAS* (fig. S5C). We validated that cGAS D319A was enzymatically inactive, as it failed to produce cGAMP after stimulation with herring testis DNA (Fig. 4E). Nevertheless, we recorded a similar elevation of ISG expression in ATRX mutant cells expressing D319A mutant or WT cGAS (Fig. 4F), demonstrating that the enzymatic activity of cGAS is not required to mediate the interferon response observed in the context of loss of ATRX.

To also explore the role of STING in cGAS-mediated interferon activation induced by ATRX loss of function, we generated BJ-hTERT fibroblasts KO for *STING1* (fig. S5D) and confirmed that cGAS and MAVS remained functional (fig. S5E-G). We then engineered clones KI or KO for *ATRX* on a *STING1* KO background and assessed their transcriptional status. Despite the absence of STING, we observed a significant elevation of interferon signalling in ATRX deficient cells (Fig. 4G, fig. S5H), with the changes in hallmark pathways recapitulating those observed in ATRX mutant cells on a *STING1* WT background (Fig. 4H, 2H). Finally, when we knocked-down ATRX in THP-1 cells KO for *IRF3*, a key transcription factor triggered downstream of cGAS-dependent interferon induction after DNA sensing, we observed an interferon response similar to that seen in WT THP-1 cells (fig. S5I-J).

Overall, these data suggest that, although the induction of interferon in ATRX deficiency involves cGAS, this induction is not mediated through the cGAS-dependent canonical DNA sensing pathway.

### Y1758C ATRX retains its capacity to bind chromatin

Considering the role of ATRX as a master genomic regulator^36^, and the finding that transcriptional activation of interferon signalling genes is independent of classical sensing pathways in the context of ATRX deficiency, we next analysed chromatin alterations resulting from a loss of function of ATRX. First, we interrogated the chromatin distribution of KI *versus* WT ATRX. In BJ-hTERT fibroblasts WT or KO for *CGAS*, both WT and KI mutant ATRX were extracted from nuclei at low salt concentration, indicating a similar degree of association to chromatin (Fig. 5A, fig. S6A). We then used Cleavage Under Targets & Tagmentation (CUT&Tag)^37^ to characterize the genomic distribution of ATRX in BJ-hTERT fibroblasts WT and KI for *ATRX*, WT or KO for *CGAS*. This analysis revealed that the Y1758C mutation did not alter ATRX binding to chromatin (with 15,475 of 15,494 individual peaks unchanged) (Fig. 5B), and that cGAS was not involved in determining the localization of ATRX on chromatin (fig. S6B). Focusing next on ATRX-bound regions shared between biological replicates, we found that a large fraction of WT ATRX occupied promoters (TSS < 3 kb), localising near 2042 genes (Fig. 5C). Notably, analysis of our RNA-seq data showed that the expression of a subset of between 20 - 24% of these ATRX-associated genes exhibited a statistically significant down or upregulation in *ATRX* KI or KO cells, as compared to *ATRX* WT cells (Fig. 5D-E).

**Figure 5.**
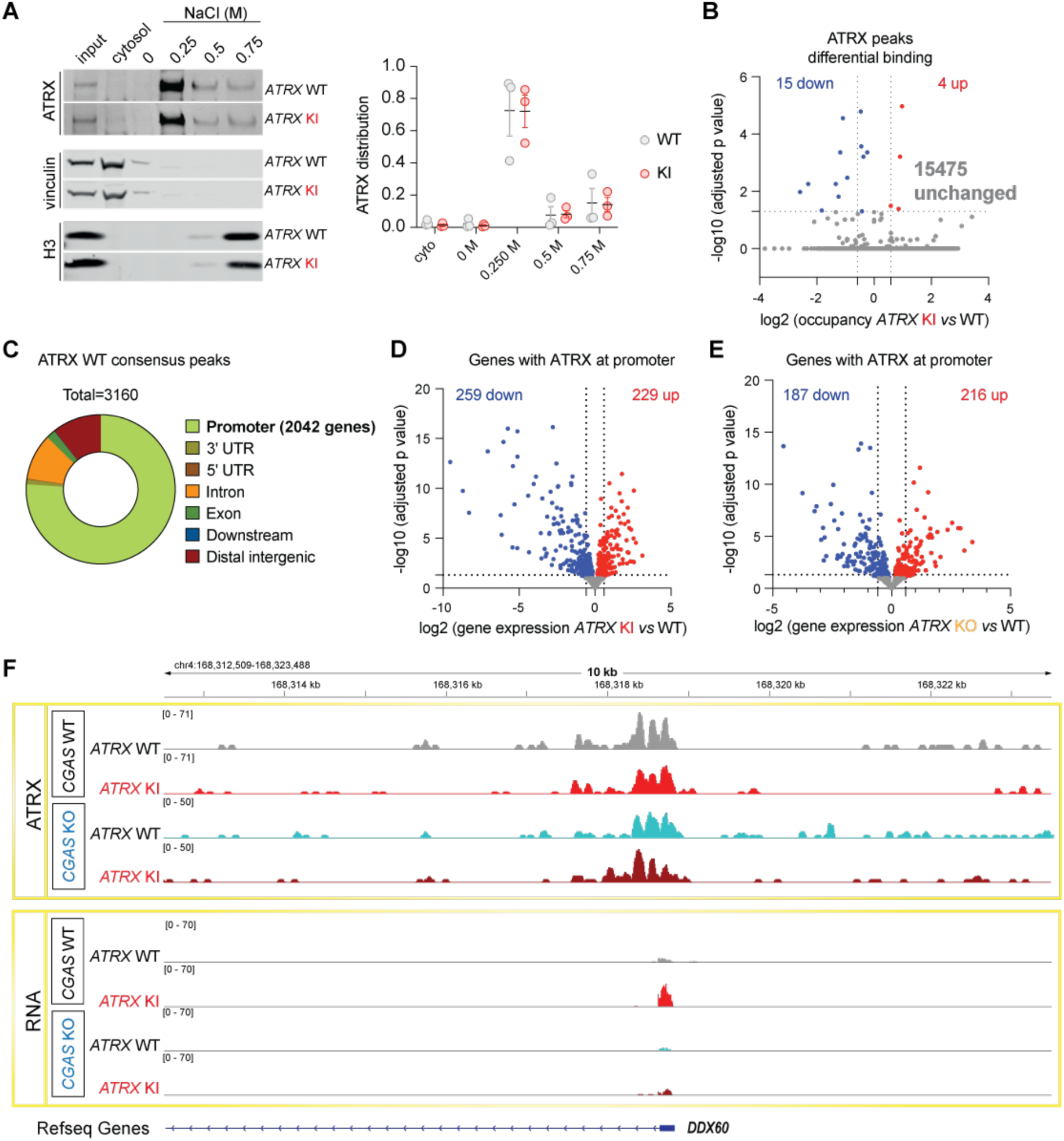
ATRX Y1758C retains its chromatin binding capacity. (A) Subcellular fractionation of BJ-hTERT fibroblasts WT or KI for *ATRX*. Cells were separated into cytosolic and nuclear fractions, and nuclear pellets subjected to sequential elution with the indicated concentrations of NaCl. Amounts of ATRX and indicated control proteins (vinculin and H3) were assessed by western blot (left); Quantification of the distribution of WT and KI ATRX in the different fractions (right). The band intensities of ATRX in the cytosol and in each of the 0, 0.25, 0.5, 0.75 M salt fractions were summed and set to 1, and the intensity of each fraction then expressed as a proportion of that sum [three independent experiments, mean +/-SEM]. (B) Genome-wide differential binding analysis of ATRX between KI and WT *ATRX* cells, individual peaks of two biological replicates are considered for each genotype; grey, red and blue dots represent peaks that are, respectively, unchanged, enriched or depleted in ATRX KI cells (adjusted p value <; 0.1). (C) Genomic distribution of WT ATRX consensus peaks based on their position in relation to genes. Consensus peaks were defined as peaks shared between biological replicates. (D-E) Volcano plots depicting differential expression of genes whose promoters are occupied by WT ATRX. Expression was assessed by RNA-seq and compared between *ATRX* KI (D) or KO (E) and WT cells; red and blue dots represent up- and down-regulated genes respectively (adjusted p value <; 0.1). (F) Representative genome browser tracks showing levels of ATRX and RNA at the upregulated gene *B2M* in BJ-hTERT fibroblasts WT or KI for *ATRX*, and WT or KO for *CGAS*.

Overall, these data show that, while the Y1758C mutation in ATRX does not affect its binding to chromatin, this mutation does result in changes in the transcription of genes in the vicinity of ATRX bound sites (Fig. 5F).

### Loss of function of ATRX alters the nucleosomal landscape at genes in a cGAS-dependent manner

We hypothesized that altered transcriptional regulation of genes might be associated with changes in the chromatin landscape in cells lacking functional ATRX, particularly focusing on the role of the histone chaperone DAXX and histone variant H3.3 in transcription^38, 39, 40, 41^. We employed CUT&Tag to assess the levels of DAXX and H3.3 at all genes differentially expressed in cells deficient for ATRX function (Fig. 2G). Importantly, while DAXX occupancy correlated variably overall with differentially expressed (down and upregulated) genes (Fig. 6A, fig. S7A), we observed a preferential reduction of DAXX levels at upregulated ISGs (Fig. 6A, fig. S7B). In contrast, changes in H3.3 levels correlated positively with transcriptionally upregulated genes in ATRX deficient cells (Fig. 6B, 6C, fig. S7C). To further investigate the link between DAXX and ISG expression, we depleted DAXX in ATRX deficient cells (fig. S7D), finding that cells mutated for *ATRX* and heterozygous KO for *DAXX* demonstrated an exaggerated interferon response, as reflected by the enhanced expression of *IFNB1* and ISGs (Fig. 6D-E) (with cells completely deficient for DAXX being non-viable).

**Figure 6.**
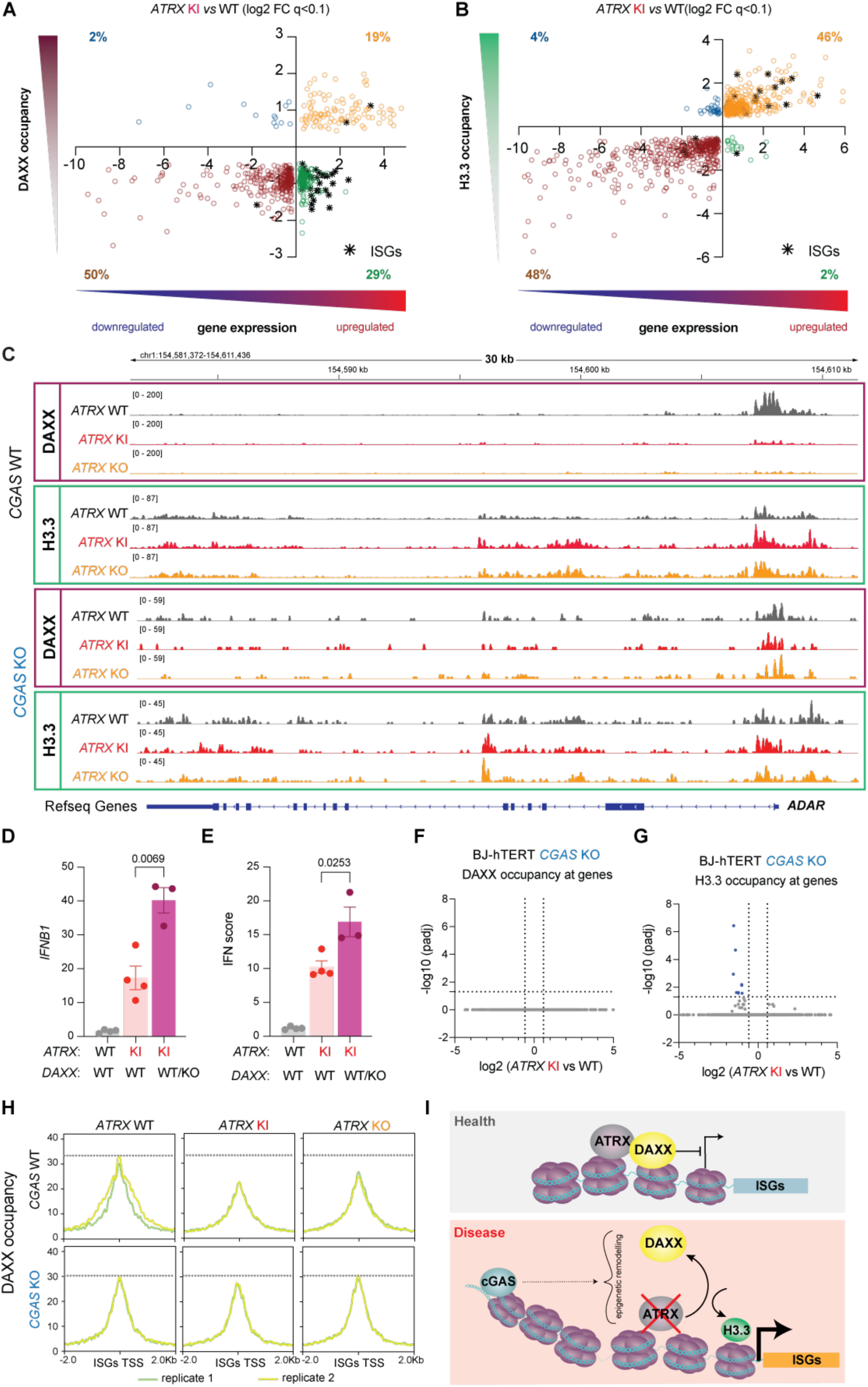
Loss of ATRX function provokes DAXX depletion and H3.3 enrichment at interferon stimulated genes in a cGAS-dependent manner. (A-B) Dot plots of genes exhibiting significant (adjusted p value (q) <; 0.1, x-axis) changes (‘FC’ signifies fold change) in both transcription and in DAXX (A) or H3.3 (B) levels in *ATRX* KI compared to WT cells. Quadrant colour coding is as follows: blue = genes enriched in DAXX or H3.3 and downregulated in expression; orange = genes enriched in DAXX or H3.3 and upregulated in expression; brown = genes depleted of DAXX or H3.3 and downregulated in expression; green = genes depleted of DAXX or H3.3 and upregulated in expression; ISGs are denoted separately by a black asterisk, while other genes are represented by empty circles. (C) Representative genome browser tracks showing levels of H3.3, DAXX at the ISG *STAT1*. (D-E) Relative expression of *IFNB1* (D) and interferon (IFN) score (median of the relative expression of the following ISGs: *IFI27, IFI6, STAT1, IFIT2, IRF7, IRF9, IFIH1, MX1*) (E) measured by qPCR in BJ-hTERT fibroblasts WT or KI for *ATRX*, and WT or heterozygous KO for *DAXX* [three to four independent clones repeated three times, Kruskal-Wallis test, two-stage linear step-up procedure of Benjamin, Krieger and Yekutieli, mean +/-SEM, q values are indicated on graphs]. (F-G) Comparison of DAXX (F) and H3.3 (G) occupancy at gene bodies between BJ-hTERT fibroblasts WT and KI for *ATRX* on a *CGAS* KO background; grey dots indicate genes exhibiting no difference in DAXX or H3.3 occupancy; blue dots indicate genes with significantly altered (adjusted p value <; 0.1) H3.3 occupancy in *ATRX* KI cells as compared to WT cells. (H) DAXX occupancy assessed by CUT&Tag at +/-2 kb of transcription starting sites (TSS) of a curated list of 444 canonical ISGs in BJ-hTERT fibroblasts WT, KI or KO for *ATRX* (from left to right), and WT or KO for *CGAS* (Top and bottom). (I) In homeostasis (top), ATRX and DAXX bind near the promoters of ISGs to regulate their expression. In ATRX loss of function (bottom), DAXX is depleted, and H3.3 enriched, at the promoters of ISGs. cGAS, which remains normally bound to nucleosomes, mediates these alterations in nucleosomal patterning and subsequent changes in gene expression.

Given the cGAS-dependency of transcriptional changes mediated by the loss of function of ATRX, we next explored the behaviour of DAXX and H3.3 in cells KO for *CGAS*. Consistent with the minimal transcriptional variation observed between cGAS null cells WT or mutant for ATRX (fig. S4F), in an analysis of all protein-coding genes, DAXX and H3.3 occupancy was similar in cells mutated or WT for *ATRX* on a *cGAS* KO background (Fig. 6F-G, fig. S7E-F). Notably, at a curated list of 444 canonical ISGs, while DAXX occupancy was reduced at promotors in ATRX deficient cells when cGAS was present, this was not the case when cGAS was absent (Fig. 6H).

Collectively, these findings indicate that ATRX deficiency leads to a depletion of DAXX and enrichment of H3.3 at activated ISGs, with cGAS playing an essential role in mediating these changes in chromatin landscape and gene expression.

## DISCUSSION

In this study we demonstrate that loss of function mutations in the chromatin remodeller ATRX can result in enhanced type I interferon signalling, thereby defining a novel type I interferonopathy state. We show that while this enhanced interferon signalling is cGAS-dependent, it does not involve the canonical pathway downstream of cGAS where the production of cGAMP leads to STING activation. Rather, in the context of a loss of ATRX activity, our study reveals a non-canonical role of cGAS in regulating the genomic occupancy of DAXX and H3.3, and thus gene transcription.

The intracranial calcification seen in P1 and P2 has not been reported previously in ATR-X syndrome. However, there have been previous descriptions of cerebral white matter disease in ATR-X syndrome ^21, 22^ and recent data implicating ATRX in myelination^42^, raising the possibility that neuroinflammation may be more common in this disorder than is currently appreciated. In support of this, we observed increased ISG expression in fibroblasts derived from four of the six patients tested. Such observed variability between patients may be due to the causative mutation, or to genetic background. Certainly, the latter is the case in regard to the severity of alpha-thalassemia seen in some patients with ATR-X syndrome (quantified by the percentage of red blood cells with haemoglobin H inclusions), where manifestation depends on the length of the G-rich tandem repeat at the alpha-globin locus^27, 43^. We hypothesise that variability in the inflammatory phenomenon we report here reflects the recognised stochastic nature of perturbed chromatin accessibility and gene expression^2^. Moreover, the immunogenicity of ATRX deficiency is apparently cell type specific; thus, while *ATRX* null sarcomas demonstrate impaired adaptive and innate immune signalling after ionizing radiation^44^, *ATRX* mutations mediate an immunogenic phenotype with macrophage infiltration in neuroblastoma^45^.

We show that the cellular consequences of the Y1758C mutation reported here are largely equivalent to the loss of ATRX. However, we demonstrate that Y1758C ATRX protein is stable and binds normally to chromatin. How this mutation affects ATRX activity requires further study. It is unlikely that this mutation directly disrupts the interaction between ATRX and DAXX, as the tyrosine at 1758 falls outside the DAXX-Binding Domain of ATRX (amino acids 1260-1289)^46^. More probable, we hypothesise that the Y1758C substitution impairs the ATPase/helicase activity of ATRX. ATR-X syndrome-associated mutations are considered to act as hypomorphs, due to reduced protein level and/or through a disruption of protein function. Of note, three previously described missense substitutions in ATRX (V1552F, K1650N and L1746S)^29^ were shown to result in a relatively stable mutant protein, and found to impact chromatin remodelling activity. We speculate that the Y1758C substitution might also affect the helicase activity of ATRX, and thereby its ability to remodel chromatin function (figure S8).

It is known that chromatin remodelling complexes, such as ATRX, can influence the association of transcription regulators with chromatin, thereby redefining the chromatin landscape and gene transcription^47, 48^. Here, we show that ATRX deficiency impaired nucleosomes composition at transcriptionally dysregulated genes. Specifically, we observed reduced levels of DAXX and enrichment of H3.3 at upregulated ISGs. Our findings support the notion that the loss of ATRX and DAXX has a cooperative and/or synergistic effect in activating an interferon signalling transcriptional program, as DAXX depletion enhances the interferon response in ATRX deficient cells.

In the context of ATRX deficiency, it seems unlikely that DAXX is the chaperone responsible for H3.3 deposition. It is known that H3.3 is present at both actively transcribed genes and constitutive heterochromatin, with its deposition regulated by distinct mechanisms in these different genomic contexts^38^. ATRX/DAXX mediates H3.3 deposition at regions enriched with H3K9me3, whereas the HIRA complex can deposit H3.3 in transcriptionally active genomic regions such as gene bodies and promoters^49, 50^. Elucidating the pathways involved in the altered dynamics of DAXX and H3.3 at chromatin, and determining if these changes work alongside the recruitment of STATs/IRFs to boost the expression of inflammatory genes, will be necessary to understand the regulatory mechanisms underpinning the autoinflammation that we report in the context of ATRX deficiency.

In the absence of cGAS, ATRX deficient cells demonstrate behaviour equivalent to ATRX WT cells, indicating that cGAS plays a central role in the chromatin alterations associated with ATRX deficiency. While an important function of the interaction of cGAS with nucleosomes is to prevent its autoactivation through sequestration away from cellular DNA^51, 52^, sequential salt elution showed that cGAS distribution on chromatin remained consistent between WT and ATRX mutant cells (Fig. 4D), suggesting that ATRX does not affect the ability of cGAS to bind to nucleosomes. Rather, our study indicates a non-canonical role for nuclear cGAS in regulating transcription in the context of ATRX deficiency, independent of the activation state of cGAS. This role thus contrasts with our previous observation of disrupted histone stoichiometry leading to cGAS-dependent, cGAMP-mediated enhancement of interferon signalling^53^, or the finding that BAF can outcompete cGAS on DNA^54^, both of which serve to limit the activation of cGAS by cellular DNA. Thus, future research should explore how cGAS contributes to changes in chromatin remodelling consequent upon ATRX deficiency.

In summary, our findings establish a model where the loss of ATRX function affects chromatin landscape, leading to the altered distributions of DAXX and H3.3, and associated changes in gene transcription. In this context, cGAS exerts a non-canonical role in regulating chromatin remodelling mediated by ATRX, a process that occurs independently of the activation state of cGAS (Fig. 6I).

## MATERIALS AND METHODS

### Clinical reports

Clinical information and samples were obtained with informed consent. The study was approved by the Comité de Protection des Personnes (ID-RCB/EUDRACT: 2014-A01017-40) in France, and the Leeds (East) Research Ethics Committee (REC reference: 10/H1307/2; IRAS project ID: 62971) in the UK.

### P1 (family 1)

This patient was born to non-consanguineous parents of white European ancestry. There was no family history of note. The patient was delivered at full term demonstrating intrauterine growth retardation. There were significant early problems with feeding, poor weight gain and recurrent upper respiratory tract infections. The patient demonstrated a delay in articulation and comprehension and subsequently exhibited significant learning difficulties leading to placement in a special school. The patient required surgical correction of an umbilical hernia and underwent tibial tendon transfer. Chondromatous lesions of the distal femoral metaphyses were also identified.

The patient presented with a single seizure. Cerebral CT showed multiple small round high signal spots of calcification in the cerebellar hemispheres, middle cerebral peduncle, and pons. Supratentorially, larger, dense calcifications were present in the caudate, globus pallidus and putamen, and small dots of calcification were seen in the right frontal and left parietal white matter / grey-white matter junction. There was a left parietal cystic area surrounded by calcium deposits. On T2 weighted cerebral MRI there was left hemispheric swelling, and a high signal white matter lesion which appeared solid, with heterogeneous signal intensity and surrounding contrast enhancement. Biopsy of the left sided cerebral lesion demonstrated features of ischaemic necrosis, with numerous foci of calcification and a minor lymphocytic infiltrate of the medium-sized vascular walls in the absence of granuloma, tumour or infection.

ISG expression in blood was persistently elevated on each of 6 occasions assayed. Exome sequencing revealed a p.(Tyr1758Cys) (Y1758C, c.5273A>G) substitution in ATRX, which was inherited from one parent. An unaffected sibling and relative were found not to carry the variant.

The patient has experienced persistently poor weight gain. Investigation for the presence of haemoglobin H (HbH) inclusions was normal.

### P2 (family 2)

This patient is the only child of unrelated white British parents. There is no family history of note beyond a unilateral ptosis and strabismus in one parent. The patient was delivered prematurely, following a pregnancy complicated by early rupture of membranes and poor fetal growth. The patient required CPAP, was fed by nasogastric tube, received phototherapy for jaundice and was treated for presumed sepsis. Atrial and ventricular septal defects were identified on echocardiography, which closed spontaneously. The patient experienced mild developmental delay, three generalised febrile seizures, and mild brachycephaly with a tall forehead, a short nose, a tented upper lip and high arched palate. The patient was diagnosed with bilateral asymmetric retinal vascular disease, leading to recurrent vitreous haemorrhages, a vitrectomy, retinal detachment and loss of vision in one eye. MRI revealed numerous peripheral enhancing lesions with haemorrhagic foci in the cerebrum, putamen and left thalamus, considered most likely to represent small vessel vasculitis. A CT scan showed extensive calcification. The patient presented with headaches, and further imaging revealed progressive suspected inflammatory changes. The patient developed a left hemiparesis, and presented with symptomatic hypoalbuminemia, considered to have been precipitated by a bout of gastroenteritis. Investigations confirmed a protein losing enteropathy, with diffuse thickening and signal enhancement of the small bowel wall, several small gastric varices, and diffuse low attenuation of the liver on ultrasound suggestive of fatty infiltration in the absence of portal hypertension. The patient has continued to experience intermittent swelling of legs and genitals, and had two fractures requiring surgery.

Investigations have included a normal karyotype, and normal testing of the following genes: *DADA2*, *CTC1*, *STN1*, *SNORD118*, and the following two panels: “Inflammatory disorders” and “interferonopathy-related genes”. Exome sequencing revealed no other putative mutations beyond the Y1758C substitution in ATRX. Autoantibody screen, cardiolipin, glycoprotein 1, Beta-2-glycoprotein antibodies, rheumatoid factor, double stranded DNA, C4, C4 normal IgG and IgM, transferrin, electrolytes liver function tests, CK, thyroid function, full blood count, were all essentially normal.

ISG expression in blood was persistently elevated. Targeted Sanger sequencing confirmed a p.(Tyr1758Cys) (Y1758C, c.5273A>G) substitution in ATRX, which was inherited from one parent.

Now, the patient demonstrates autistic traits, moderate learning difficulties, and facial features consistent with ATR-X syndrome. HbH inclusions were not detected on two occasions.

### Genetic studies

DNA was extracted from whole blood using standard methods. Exome sequencing was performed on genomic DNA from patients 1 and 2, and the parents of patient 1, using a SureSelect Human All Exon kit (Agilent Technologies) for targeted enrichment, and Illumina HiSeq2000 for sequencing. The variant c.5273A>G / p.Tyr1758Cys in exon 21 of ATRX was assessed using the *in silico* programs SIFT (http://sift.jcvi.org) and Polyphen2 (http://genetics.bwh.harvard.edu/pph2/). Population allele frequencies were obtained from the gnomAD database (http://gnomad.broadinstitute.org). Confirmatory Sanger sequencing was undertaken in both patients, and used to assess the genetic status of other family members. The reference sequence used for primer design and nucleotide numbering was NM_000489.3. Primers for Sanger sequencing of ATRX exon 21: ATRX Ex21F tgtgtttttgacatgagcatttca, ATRX Ex21R tgaggctttacacctggcaa, generating a PCR product of 506bp.

### RT-qPCR quantification and interferon stimulated gene (ISG) gene expression

Whole blood was collected into PAXgene tubes (Qiagen) and total RNA extracted using a PreAnalytix RNA isolation kit. Interferon (IFN) scores were generated as previously described^55^.

For NanoString ISG analysis, total RNA was similarly extracted from whole blood with a PAXgene (PreAnalytix) RNA isolation kit. Analysis of 24 interferon stimulated genes and 3 housekeeping genes (probes of interest [n = 24]: *IFI27*, *IFI44L*, *IFIT1*, *ISG15*, *RSAD2*, *SIGLEC1*, *CMPK2*, *DDX60*, *EPSTI1*, *FBXO39*, *HERC5*, *HES4*, *IFI44*, *IFI6*, *IFIH1*, *IRF7*, *LAMP3*, *LY6E*, *MX1*, *NRIR*, *OAS1*, *OASL*, *OTOF*, and *SPATS2L*; reference probes [n = 3]: *NRDC*, *OTUD5*, and *TUBB*) was conducted using the NanoString customer designed CodeSets according to the manufacturer’s recommendations (NanoString Technologies). Data were processed with nSolver software (NanoString Technologies). The data were normalized relative to the internal positive and negative calibrators, the three reference probes, and the control samples. Scores above 2.75 are designated as positive.

In cell lines, total RNA was extracted using the RNeasy Micro Kit (Qiagen, 74004), and reverse transcription performed with the High-Capacity cDNA Reverse Transcription Kit or SuperScript™ IV VILO™ Master Mix (ThermoFisher Scientific). Levels of cDNA were quantified by RT-qPCR using TaqMan Gene Expression Assays (ThermoFisher Scientific), normalized to *HPRT1* (Hs03929096), *ACTB* (Hs01060665_g1), *VCL* (Hs00419715_m1) and *TBP* (Hs00427620_m1). Relative quantitation of target cDNA was determined using the formula 2−ΔΔCT, with ΔΔCT representing fold increase above control. Interferon (IFN) scores were calculated as the median of the fold change expression of selected ISGs: *IFI27* (Hs01086370_m1), *RSAD2* (Hs01057264_m1), *MX1* (Hs00895608_m1), *ISG15* (Hs00192713_m1), *IFI44L* (Hs01060665_g1), *IFIT1* (Hs00356631_g1), *TNFAIP3* (Hs00234713_m1), *STAT1* (Hs01013996_m1), *ALDH1A1* (Hs00946916_m1), *IFI6* (Hs00242571_m1), *IFIH1* (Hs00223420_m1), *OAS1* (Hs00973637_m1). Other probes used in this study were: *IFNB1* (Hs01077958_s1) and *MAVS* (Hs00920075).

### Cell culture

Fibroblasts and 293FT (Invitrogen, R70007) cells were grown in DMEM-Glutamax supplemented with 10% FBS and 1% P/S (10,000 U/mL). Primary fibroblasts from P1 and P2 were generated in this paper; P3 to P6 are part of the Gibbons laboratory cell bank, P3: c.666G>T / p.(W222C); P4: c.797A>G / p.(Y266C); P5: c.5039T>C / p(.I1680T); P6: c.413_414delAA / p.(K138fs); controls were purchased from Gibco and Innoprot; patient TREX1 (AGS261.1): p.(Glu20Glyfs*82)/p.(Ala81Serfs*21); patient DNASEII (AGS85.1): homozygous p.(Gly116Ala) and patient RNASEH2A (AGS070.1): p.(Arg235Gln)/p.(Arg25Arg) are part of the Crow laboratory cell bank. BJ-hTERT WT and null for cGAS were obtained from the Chen laboratory (University of Texas, Dallas)^33^; THP-1 and *IRF3* KO THP-1 cells were from ATCC (TIB-202) and InvivoGen respectively.

Fibroblast stimulations were conducted by treating cells for four hours with 1 µg/mL herring testes (HT)-DNA (D6898; Sigma-Aldrich) complexed with Lipofectamine 2000 (Invitrogen) according to the manufacturer’s instructions, with 1 µM diABZI (Cayman, 280541), or with 100 ng/mL Poly (I:C) (Invitrogen, cat nb tlrl-piclv, (HMW) / LyoVec™) for six hours.

### ISG Reporter HEK 293 Cells

Interferon Regulatory Factor (IRF)-Inducible SEAP Reporter HEK293 (HEK-Blue™ ISG) cells (InvivoGen, hkb-isg-1) were used as an interferon activity reporter. 50,000 cells were seeded per well of a 96 well-plate and stimulated with 180 μL of THP-1 cell supernatant transduced with sh-RNAs targeting *ATRX* or control sh-RNAs, and collected at day eight post-transduction. Cells were stimulated for 24 hours in a final volume of 200 μL. Two wells were tested per experimental condition. Culture supernatants were collected, and levels of embryonic alkaline phosphatase (SEAP) secreted by HEK-Blue™ ISG cells determined with QUANTI-Blue™ Solution (turning purple/blue in the presence of SEAP) calculated from the OD at 650 nm read using a microplate reader. Readings from unstimulated cells were subtracted, and measurements normalised relative to THP-1 sh-control conditions.

### Induced pluripotent stem cell (iPSC) culture and neural differentiation

The iPSC QOLG-1 cell line was obtained from Kathy Evans (Centre for Genomic and Experimental Medicine Institute of Genetics and Cancer, Edinburgh. UK), acknowledging the Wellcome Trust Sanger Institute as the source of the HipSci HPSI1113i-qolg_1 human induced pluripotent cell line (ECACC 77650145). iPSCs were made using a CytoTune iPS Reprogramming kit obtained from Life Technologies by Sanger as the ‘the buyer’, under a Limited Use Label Licence. iPSCs were maintained on vitronectin (Stem Cell Technologies, 7180), with a complete media change every day in Essential 8 media (Life Technologies, A1517001). 0.5mM EDTA in PBS was used for passaging. Accutase (Stem cell Technologies 07920) and 10 μM Rho-associated kinase (ROCK) inhibitor (Sigma-Aldrich Y0503) were used for single-cell dissociation purposes. iPSCs were collected at passage 38 to 40.

Neural progenitor cells (NPCs) were established following the STEMdiff Neural System based on STEMdiff SMADi Neural Induction kit (Stem cell Technologies, 08581). Briefly, embryoid bodies (EBs) were generated using AggreWell™800 plates, with each EB containing approximately 10,000 cells. EBs were fed daily for four days with STEMdiff Neural Induction Media + SMADi. At day five, EBs were replated on Matrigel coated dishes and fed daily for six days with STEMdiff Neural Induction Media + SMADi. Neural Rosettes were dislodged from EBs using STEMdiff Neural Rosette Selection solution (Stem cell Technologies, 05832), replated on Matrigel coated dishes, and fed daily with STEMdiff™ Neural Induction Media + SMADi. When cells reached confluency, they were split in two using Accutase, and then replated (passage 1). From passage one onwards NPCs were fed daily with STEMdiff Neural Progenitor Media (Stem cell Technologies, 05833).

NPCs were used at passage two, three or five to generate forebrain neurons following the instructions of STEMdiff Forebrain Neuron Differentiation Kit (Stem cell Technologies, 08600) and STEMdiff Forebrain Neuron Maturation Kit (Stem cell Technologies, 08605). Briefly, NPCs were plated on Poly-L-Ornithine/Laminin coated dishes and fed daily with STEMdiff forebrain differentiation media for six days. Generated neural precursors were replated on Poly-L-Ornithine/Laminin coated dishes, fed with STEMdiff forebrain neuron maturation kit twice a week, and cultured for three to four weeks.

### Knock-down and over-expression of genes

Lentiviral constructs containing shRNA targeting *ATRX* (pLKO.1_shATRXn1: TRCN0000342811, pLKO.1_shATRXn2: TRCN0000013590, pLKO.1_shATRXn3: TRCN0000352788) were from Sigma-Aldrich. pLKO.1 empty vector (sh-control1) was from Open Biosystems, and pLKO.1 control shRNAs (scramble shSCR, SHC002 (sh-control2, and non-target shNTgt, SHC016, sh-control3) were from Sigma-Aldrich. Lentiviral particles were produced by Lipofectamine 2000 transfection of 293FT cells with shRNA constructs, psPAX2, and pCMV-VSV-G. Supernatant was used to infect THP-1 cells. Two days after transduction, cells were cultured with media containing 1.5 µg/mL puromycin (Sigma-Aldrich). Cells were collected for analysis five and eight days after transduction.

Expression of *MAVS* was downregulated by using 20 nM siRNAs targeting *MAVS* (LifeTechnologies #s41746, #148763) or negative control#1 (LifeTechnologies, Silencer Select, Cat#4390843). All were delivered to cells using Lipofectamine RNAiMAX (Invitrogen) according to the manufacturer’s instructions. Cells were analysed 72 h after transfection.

For cGAS WT and D319A overexpression in BJ-hTERT fibroblasts, transfection was performed by using Lipofectamine LTX. Briefly 250,000 cells were seeded in one well of.a 6-well plate; 1:1 DNA/supplement ratio was used, and 0.82 ug of WT or 1.2 ug of D319A cGAS plasmids were complexed to 2.46 or 3.6 uL of lipofectamine, respectively. Complexes were added on cells and incubated for four hours. Supernatants were then removed and replaced with full growth media. Cells were collected for analysis three days after transfection. The plasmids used were pEFBOS-cGAS-WT-Flag-His and pEFBOS-cGAS-D319A and were kindly gifted by the laboratory of Andrea Ablasser (EPFL, Lausanne, Switzerland).

### Western blot

Proteins from whole cell lysates were extracted using RIPA lysis buffer (25 mM Tris-HCl pH 8.0, 1% IGEPAL 60, 150 mM NaCl, 1.5 mM MgCl2, 0.05 % SDS, 0.5% sodium deoxycholate, Benzonase (Sigma-Aldrich, E1014) 750 U/mL) supplemented with protease and phosphatase inhibitor tablets (ROCHE). Reducing Buffer was added to proteins and heated for 5 min at 95°C. Samples were resolved on 3-8% Tris-acetate or 4–12% Bis-Tris and then transferred to a nitrocellulose membrane using the iBlot 2 Dry Blotting System (LifeTechnologies). Membranes were blocked and primary antibodies incubated overnight at 4°C. Membranes were washed and incubated with secondary antibodies for 45 min at room temperature. Signal was detected using the Odyssey CLx System (LI-COR). Quantifications were performed using Image Studio 5.2 (LI-COR System). Antibodies and dilution used were: ATRX (NBP2-50111, 1/500); Ser172-TBK1 (CST 5483, 1/500), Tyr701-Stat1 (CST 7649, 1/700); STAT1 (CST 9176, 1/1000); Ser396-IRF3 (CST 4947, 1/700); vinculin (CST 13901, 1/1000), S366-STING (CST 19781, 1/1000), STING (Millipore MAB7169, 1/1000), NF-κB p65 (CST 6956, 1/1000), Phospho-NF-κB p65 (Ser536) (CST 3031, 1/1000), cGAS (CST 15102, 1/500), α-tubulin (Sigma-Aldrich T6074, 1/5000), Histone H3 (CST 4499, 1/5000), Histone H2B (CST 12364, 1/2000).

### Subcellular fractionation

For subcellular salt extraction fractionation, we followed a published protocol with minor changes ^51^. Cells were pelleted, washed in PBS, resuspended in extraction buffer (10 mM Hepes pH 7.9, 10 mM KCl, 1.5 mM MgCl2, 0.34 M sucrose, 10% glycerol, 0.2% IGEPAL 60), and incubated on ice for 10 minutes with occasional vortexing. Nuclei were spun at 3500 x g for 5 minutes at 4° C. The cytosolic fraction (supernatant) was collected for further analysis. Nuclei were then washed for one min on ice in extraction buffer without IGEPAL 60 and spun at 3500 x g for 5 minutes at 4°C. Pelleted nuclei were resuspended in zero salt buffer (3 mM EDTA, 0.2 mM EGTA), vortexed 10 seconds on/10 seconds off for one min, and incubated on ice for 30 minutes, vortexing for 15 seconds every 15 minutes. Lysates were spun at 6500 x g for five minutes at 4° C. The zero-salt supernatant was collected for further analysis. The remaining pellets were then resuspended in first salt buffer (50 mM Tris-HCl, pH 8.0, 0.05% IGEPAL 60, 250 mM NaCl), incubated on ice for 15 min with vortexing twice. Lysates were spun at 17,000 g at 4°C for 5 minutes. Supernatants were collected for further analysis. Likewise, 500 and 750 mM NaCl extractions were performed. Lastly, pellets were resuspended in 1 M NaCl salt buffer supplemented with DNase I (Invitrogen, AM2238) and 10 mM MgCl2, incubated 30 min at 37C, spun at 17,000 g at 4°C for five minutes and supernatants collected. All buffers were supplemented with complete Protease Inhibitor Cocktail (Roche). All samples were analysed by Western blots following the protocol described above.

### Immunofluorescence staining and images analysis

Cells were grown on glass coverslips in a 24-well plate and fixed with 4% paraformaldehyde in PBS for 15 min, treated with 0.1 M glycine in PBS for 20 min, permeabilized with 0.2% Triton X-100 in PBS, followed by blocking in 5% normal goat serum in PBS for one hour. Cells were incubated with primary antibody in 2% bovine serum albumin (BSA) in PBS, and incubated overnight in a humidified chamber at 4°C (MAP2, Abcam ab5392, 1/2000); NeuN (Millipore MAB377, 1/20); OCT4 (Invitrogen MA1-104, 1/500); Pax6 (Biolegend 901302, 1/100); Nestin (Novus Biological NB300266, 1/500). Coverslips were washed and incubated for one hour at room temperature with Alexa Fluor secondary antibodies (Life Technologies, 1/200). Slides were mounted using ProLong Gold Antifade Mountant with DAPI. Nikon A1R+ confocal microscope was used to acquire images analysed by ImageJ Fiji^56^.

### cGAMP ELISA

For cGAMP quantification, cells were harvested and lysed in RIPA lysis buffer ((50 mM Tris, 150 mM NaCl, 1% sodium deoxycholate, 0.03% SDS, 0.005% Triton X-100, 5 mM EDTA and complete Protease Inhibitor Cocktail (Roche)) for 30 min on ice. Lysed cells were centrifuged for 10 min at 17,000g and 4°C. The supernatant (50 μL*2) was used for the 2′3′-cGAMP ELISA assay (Cayman, 501700) according to the manufacturer’s instructions. Protein concentration in the supernatant was measured using BCA Pierce Protein assay kit and used to normalize 2′3′-cGAMP levels.

### CRISPR/Cas9

For genome editing, direct delivery of the CRISPR/Cas9 system as a ribonucleoprotein (RNP) complex consisting of Cas9 protein and single guide RNA (sgRNA) was used. For *ATRX*: A single Alt-R CRISPR-Cas9 sgRNA targeting *ATRX*, containing the target-specific crRNA region (TTCTCTCTTAGATCATTGTA) and the Cas9-interacting tracrRNA region was used. To introduce the Y1758C mutation by homologous recombination, CRISPR-Cas9 was used with a single-stranded oligodeoxynucleotide (ssODN) of 200 bp (CTAATGCATGTGTTTTTGACATGAGCATTTCATTGGGGAATTTTTCTAAGTCTTTTTAACGATTGTTAATGTTGTTAATGAATAATGTCTTCTCTCTTAGGTCACTGCATGGTTAATTTTATCAAGGAAAATTTACTTGGATCCATTAAGGAGTTCAGGAATAGATTTATAAATCCAATTCAAAATGGTCAGTGTGCAGA). SgRNA, ssODN and Alt-R S.p. Cas9 Nuclease 3NLS (IDT 1081059) were purchased from IDT. RNP complex, with or without ssODN, was delivered to cells by electroporation using 4D-Nucleofector system. Clonal expansion was assessed after two to four weeks, and established single clones genotyped for ATRX indels and knock-ins. PCR primers: TTCACAAGATAGCCTCACTGACC and GGTAGAATCTGCACACTGACCA. For *STING1*: the target-specific crRNA region was GCTGGGACTGCTGTTAAACG, and PCR primers AGGGGAACTGGGAGGAAGTGCC and CCAAGGCTTCCCAACCTGCTCC. For *DAXX*, the target-specific crRNA region was GCTCACATAGTCGCCCAAAG, and PCR primers CTGTTTTTGGCCTCGGCGGAGT and GCATCCCACCAACCCCCTACCT. Genome editing was assessed in pool cells prior to seeding for single-cell clonal expansion, using the ICE CRISPR Analysis Tool.

### Pathway enrichment analysis

GSEA^57^ was carried out using hallmark gene sets from MSigDB^58^ and recommended settings (parameters scoring_scheme classic and metric Signal2Noise). For fig. S7B, Enrichr tool^59^ was used.

### RNA Sequencing and analysis

For iPSCs and neural lines, three independent clones per genotype and one technical replicate were used. RNA was extracted using RNeasy Micro Kit (Qiagen, 74004). RNA samples were assessed using the Fragment Analyser Automated Capillary Electrophoresis system (Agilent Technologies Inc, 5300) with the Standard Sensitivity RNA Analysis kit (DNF-471-0500) for quality and integrity of total RNA, and then quantified using the Qubit 2.0 Fluorometer (Thermo Fisher Scientific Inc, Q32866) with the Qubit RNA HS Assay kit (Q32855). Libraries were generated using the QuantSeq 3’mRNA-Seq Library Prep kit (FWD) protocol for Illumina (Lexogen Inc, #015). Single read (1×75bp) sequencing was performed on the NextSeq 550 platform (Illumina Inc, SY-415-1002) using the NextSeq 500/550 High-Output v2.5 kit (20024906). Libraries were combined in a single equimolar sample based on Qubit and Bioanalyser assay results, and run across a High Output v2.5 Flow Cell. The QuantSeq and differential data analyses were performed following pipeline integration on the Bluebee genomics analysis platform.

For anifrolumab treatment, three independent clones per genotype were used and cells were treated for 24 hours with 10 μg/mL of anifrolumab (MedChemExpress HY-199168) or water. Total RNA was extracted and processed as previously described to generate libraries using the QuantSeq 3’mRNA-Seq Library Prep kit (FWD) protocol for Illumina. As an initial processing step, the raw fastq files were pre-trimmed from the 5’ end of the first poly-A sequence of length 10 or more. This trimming removed the poly-A sequences themselves along with any downstream nucleotides. The trimming was performed using a custom script implemented in Biopython^60^. Read counts over human genes were then obtained by running the pre-trimmed fastq files through the nf-core RNA-Seq pipeline using fastp^61^ for adapter trimming, STAR^62^ for alignment, and Salmon^63^ for quantification. The gene level counts obtained were then used to perform pairwise differential expression analyses using DESeq2^64^. The code used to process the data and plot the results is available at https://github.com/hyweldd/2025_atrx_quantseq_pipeline.

For BJ-hTERT lines, three independent clones per genotype and one technical replicate were used. Libraries were prepared using the NEBNEXT Ultra II Directional RNA Library Prep kit (NEB #7760), and the Poly-A mRNA magnetic isolation module (NEB #E7490) used according to the manufacturer’s protocol. Final libraries had an average peak size of 286 bp. Fragment size and quantity measurements were used to calculate molarity for each library pool. Sequencing was performed on the NextSeq 2000 platform (Illumina Inc, #20038897) using NextSeq 2000 P3 Reagents (100 Cycles) (#20040559).

RNA-Seq read data were processed through the nf-core/rnaseq (v3.10.1) pipeline (doi: 10.5281/zenodo.1400710) using genome reference, GRCh38 (GRCh38.p13) and Ensembl gene models. We used the DESeq2 package (version 1.42.1) in R (version 4.3.3) to perform differential gene expression analysis between various genotypes^64^. We obtained gene counts from the nf-core/rnaseq pipeline output^65^ (“salmon.merged.gene_counts_length_scaled.tsv”) and imported them into DESeq2 using the DESeqDataSetFromMatrix method. We normalized the counts using the variance stabilizing transformation (VST) and performed principal component analysis (PCA) to visualize the sample clustering. DESeq2 fitted a negative binomial generalized linear model with genotype as the main factor and used the Wald test to identify differentially expressed genes. We applied the ashr method to shrink the log2 fold changes for estimation stability and improved ranking of genes^66^. We also calculated s-values, which are a transformation of p-values that control the false discovery rate (FDR) while incorporating prior information on effect sizes.

For transcriptomic assays, GSEA^57^ was carried out using hallmark gene sets from MSigDB^58^ and recommended settings (scoring_scheme classic and metric Signal2Noise).

### CUT&Tag processing and data analysis

Two independent clones and two technical replicates were used per genotype. Experiments were performed following the protocol described in^37^. Briefly, fresh cells (100,000 cells per reaction) were harvested and lysed. Nuclei were isolated and bound to Concanavalin A (ConA)-coated beads. Primary antibodies were: mAb IgG XP Isotype Control CST 66362, ATRX rabbit ab97508, DAXX Prestige-HPA065779, H3.3 Epicypher 13-0061. 0.5ug of antibody in antibody buffer (0.01% BSA in wash buffer) was added to beads and incubated for two hours at room temperature, then overnight at 4°C. Beads were cleared, mixed with the secondary antibody (Guinea Pig anti-Rabbit IgG, antibodies-online ABIN101961) at a dilution of 1:100 in antibody buffer, and incubated for one hour at room temperature. Beads were washed and resuspended in Protein A/G-Tn5 (Epicypher 15-1017) for two hours. Tagmentation was performed for 1 h at 37°C in 300-Wash buffer supplemented with 10 mM MgCl2. Beads were washed and tagmented fragments released. Following 1 h incubation at 58°C, SDS was neutralized with 0.67% Triton-X100. Barcoded i5 (v2_Ad1) and i7 (v2_Ad2) primer^67^ and NEBNext 2x Master mix (NEB, M0541L) were added, and tagmented fragments amplified by PCR. PCR clean-up was performed using AMPure XP bead slurry following the manufacturer’s instructions. Equimolar libraries were made, and sequencing (2×50) was performed on the NextSeq 2000 platform using the NextSeq 1000/2000 P2 Reagents (100 cycles) v3 Kit (#20046811).

Sequencing reads were processed with the nf-core/cutandrun pipeline (3.2.2) (doi: 10.5281/zenodo.10606804), implemented in Nextflow (v24.04.2). Briefly, Adapter and quality trimming of reads was performed using Trim galore (0.6.6). Reads were aligned using Bowtie2 (2.4.4) to the human reference genome (GRCh38.p13) and to an E. coli K-12 MG1655 (spike-in). Both duplicate reads (picard: 3.1.0) and reads aligning within ENCODE blacklist regions (hg38 blacklist v1) were removed prior to peak calling. Peaks were called with SEACR (1.3) (stringent,threshold = 0.01). Using Galaxy^68^, 1/ bigwig files were generated using the bamCoverage tool, normalized to 1x sequencing depth (Reads Per Genomic Content (RPGC)), 2/ Peaks were annotated using Chipseeker, 3/ Diffbind tool was employed to identify differentially bound sites between two groups, and 4/ computeMatrix and PlotHeatmap tools were used to generate enrichment profiles.

For differential binding and principal component analysis (PCA), peaks from the two biological replicates (R1 and R2) were intersected using “bedtools intersect” (version v2.31.1). The union of peaks of all genotypes, using “bedtools merge”, was made to create a combined list of positions. Reads mapped at to these coordinates were counted using featureCounts (from the subread package, version v2.0.8). DESeq2 package (version 1.34.0) in R (version 4.1.2) was used to assess differential peak coverage between various genotypes^64^. The counts were normalised using the variance stabilizing transformation (VST) and performed PCA to visualize the sample clustering. DESeq2 fitted a negative binomial generalized linear model with genotype as the main factor and used the Wald test to identify differentially bound regions.

For differential binding on genes bodies, additional read counts were obtained over entire gene bodies using positions of UniprotKB/Swiss-Prot genes, using Biomart (Ensembl version 113), adding 3 kb on each side of the gene position. Again, we used the DESeq2 package (version 1.34.0) in R (version 4.1.2) to perform differential gene body coverage analysis between various genotypes.

### Quantification and statistical analysis

Statistical analyses for all experiments, except sequencing data, were performed in GraphPad Prism v9 or v10. Statistical parameters are reported in the figure legends.

### Data availability

Sequencing data generated for this study will be uploaded on GEO.

## SUPPLEMENTARY MATERIALS

Figures S1 to S8

## ACKNOWLEDGEMENTS

We thank the patients and their families for participation in this study. We thank James Chen (University of Texas at Dallas) for the kind gift of BJ-hTERT fibroblasts, and acknowledge Kathy Evans and the Wellcome Trust Sanger Institute for access to the human induced pluripotent cell line which was generated under the Human Induced Pluripotent Stem Cell Initiative funded by a grant from the Wellcome Trust and Medical Research Council, supported by the Wellcome Trust (WT098051) and the NIH/Wellcome Trust Clinical Research Facility. We gratefully acknowledge practical and intellectual help and advice all the members of the Crow laboratory. We thank Nelly Olova for advice relating to sequencing library preparation. We also wish to thank Wendy Bickmore and Andrew Jackson for critical reading of the manuscript. Funding: Y.J.C. acknowledges the European Research Council (786142-E-T1IFNs), and a state subsidy managed by the National Research Agency (France) under the ‘Investments for the Future’ programme bearing the reference ANR-10-IAHU-01. P.R. is staff scientist at the Centre National de la Recherche Scientifique (CNRS). Y.J.C. is supported by a UK Medical Research Council Human Genetics Unit core grant (MC_UU_00035/11). R.J.G. was supported by the Medical Research Council (grant number MC_UU_12025/ unit programme MC_UU_12009/3). R.T. is a Chancellors Fellow.

## Author contributions

M.E., E.W., G.Z., A.F, T.T., C.U., N.B, E.N., S.G., M.P.R, G.I.R., L.S., R.C., A.F., J.A.M. and P.R. performed experiments; R.T., A.L and P.R were involved in experimental planning and discussion of the results, G.G. and P.G., H.D. and S.D. performed bioinformatic analyses; J.H.L., E.W., S.N., J.V., B.I., R.J.G., and Y.J.C. ascertained and characterised clinical cases; M.E. and Y.J.C. wrote and edited the manuscript with critical review by R.T., A.L, P.R. and R.J.G.

## REFERENCES

1. Clynes, D., Higgs, D.R. & Gibbons, R.J. The chromatin remodeller ATRX: a repeat offender in human disease. Trends Biochem Sci 38, 461–466 (2013).

2. Truch, J. et al. The chromatin remodeller ATRX facilitates diverse nuclear processes, in a stochastic manner, in both heterochromatin and euchromatin. Nat Commun 13, 3485 (2022).

3. Lewis, P.W., Elsaesser, S.J., Noh, K.M., Stadler, S.C. & Allis, C.D. Daxx is an H3.3-specific histone chaperone and cooperates with ATRX in replication-independent chromatin assembly at telomeres. Proc Natl Acad Sci U S A 107, 14075–14080 (2010).

4. Valenzuela, M., Amato, R., Sgura, A., Antoccia, A. & Berardinelli, F. The Multiple Facets of ATRX Protein. Cancers (Basel) 13 (2021).

5. Gibbons, R.J., Picketts, D.J., Villard, L. & Higgs, D.R. Mutations in a putative global transcriptional regulator cause X-linked mental retardation with alpha-thalassemia (ATR-X syndrome). Cell 80, 837–845 (1995).

6. Leon, N.Y. & Harley, V.R. ATR-X syndrome: genetics, clinical spectrum, and management. Hum Genet 140, 1625–1634 (2021).

7. Bhargava, R., Lynskey, M.L. & O’Sullivan, R.J. New twists to the ALTernative endings at telomeres. DNA Repair (Amst) 115, 103342 (2022).

8. Lovejoy, C.A. et al. Loss of ATRX, genome instability, and an altered DNA damage response are hallmarks of the alternative lengthening of telomeres pathway. PLoS Genet 8, e1002772 (2012).

9. Jiao, Y. et al. Frequent ATRX, CIC, FUBP1 and IDH1 mutations refine the classification of malignant gliomas. Oncotarget 3, 709–722 (2012).

10. Hariharan, S. et al. Interplay between ATRX and IDH1 mutations governs innate immune responses in diffuse gliomas. Nat Commun 15, 730 (2024).

11. Cabral, J.M., Oh, H.S. & Knipe, D.M. ATRX promotes maintenance of herpes simplex virus heterochromatin during chromatin stress. Elife 7 (2018).

12. Lazear, H.M., Schoggins, J.W. & Diamond, M.S. Shared and Distinct Functions of Type I and Type III Interferons. Immunity 50, 907–923 (2019).

13. Crow, Y.J. & Stetson, D.B. The type I interferonopathies: 10 years on. Nat Rev Immunol 22, 471–483 (2022).

14. Ablasser, A. & Chen, Z.J. cGAS in action: Expanding roles in immunity and inflammation. Science 363 (2019).

15. Balka, K.R. & De Nardo, D. Molecular and spatial mechanisms governing STING signalling. FEBS J 288, 5504–5529 (2021).

16. Stetson, D.B., Ko, J.S., Heidmann, T. & Medzhitov, R. Trex1 prevents cell-intrinsic initiation of autoimmunity. Cell 134, 587–598 (2008).

17. Bai, J. & Liu, F. Nuclear cGAS: sequestration and beyond. Protein Cell 13, 90–101 (2022).

18. de Oliveira Mann, C.C. & Hopfner, K.P. Nuclear cGAS: guard or prisoner? EMBO J 40, e108293 (2021).

19. Aref-Eshghi, E. et al. Evaluation of DNA Methylation Episignatures for Diagnosis and Phenotype Correlations in 42 Mendelian Neurodevelopmental Disorders. Am J Hum Genet 106, 356–370 (2020).

20. Crow, Y.J. CNS disease associated with enhanced type I interferon signalling. Lancet Neurol 23, 1158–1168 (2024).

21. Lee, J.S. et al. Alpha-thalassemia X-linked intellectual disability syndrome identified by whole exome sequencing in two boys with white matter changes and developmental retardation. Gene 569, 318–322 (2015).

22. Wada, T. et al. Neuroradiologic features in X-linked alpha-thalassemia/mental retardation syndrome. AJNR Am J Neuroradiol 34, 2034–2038 (2013).

23. Deneault, E. et al. Complete Disruption of Autism-Susceptibility Genes by Gene Editing Predominantly Reduces Functional Connectivity of Isogenic Human Neurons. Stem Cell Reports 11, 1211–1225 (2018).

24. Danussi, C. et al. Atrx inactivation drives disease-defining phenotypes in glioma cells of origin through global epigenomic remodeling. Nat Commun 9, 1057 (2018).

25. Levy, M.A., Fernandes, A.D., Tremblay, D.C., Seah, C. & Berube, N.G. The SWI/SNF protein ATRX co-regulates pseudoautosomal genes that have translocated to autosomes in the mouse genome. BMC Genomics 9, 468 (2008).

26. Garrick, D. et al. Loss of Atrx affects trophoblast development and the pattern of X-inactivation in extraembryonic tissues. PLoS Genet 2, e58 (2006).

27. Gibbons, R.J. et al. Mutations in the chromatin-associated protein ATRX. Hum Mutat 29, 796–802 (2008).

28. Argentaro, A. et al. Structural consequences of disease-causing mutations in the ATRX-DNMT3-DNMT3L (ADD) domain of the chromatin-associated protein ATRX. Proc Natl Acad Sci U S A 104, 11939–11944 (2007).

29. Mitson, M., Kelley, L.A., Sternberg, M.J., Higgs, D.R. & Gibbons, R.J. Functional significance of mutations in the Snf2 domain of ATRX. Hum Mol Genet 20, 2603–2610 (2011).

30. Chen, Q., Sun, L. & Chen, Z.J. Regulation and function of the cGAS-STING pathway of cytosolic DNA sensing. Nat Immunol 17, 1142–1149 (2016).

31. Streicher, F. & Jouvenet, N. Stimulation of Innate Immunity by Host and Viral RNAs. Trends Immunol 40, 1134–1148 (2019).

32. Li, T. et al. Phosphorylation and chromatin tethering prevent cGAS activation during mitosis. Science 371 (2021).

33. Yang, H., Wang, H., Ren, J., Chen, Q. & Chen, Z.J. cGAS is essential for cellular senescence. Proc Natl Acad Sci U S A 114, E4612–E4620 (2017).

34. Sun, L., Wu, J., Du, F., Chen, X. & Chen, Z.J. Cyclic GMP-AMP synthase is a cytosolic DNA sensor that activates the type I interferon pathway. Science 339, 786–791 (2013).

35. Wu, J. et al. Cyclic GMP-AMP is an endogenous second messenger in innate immune signaling by cytosolic DNA. Science 339, 826–830 (2013).

36. Aguilera, P. & Lopez-Contreras, A.J. ATRX, a guardian of chromatin. Trends Genet 39, 505–519 (2023).

37. Henikoff, S., Henikoff, J.G., Kaya-Okur, H.S. & Ahmad, K. Efficient chromatin accessibility mapping in situ by nucleosome-tethered tagmentation. Elife 9 (2020).

38. Choi, J., Kim, T. & Cho, E.J. HIRA vs. DAXX: the two axes shaping the histone H3.3 landscape. Exp Mol Med 56, 251–263 (2024).

39. Levy, M.A., Kernohan, K.D., Jiang, Y. & Berube, N.G. ATRX promotes gene expression by facilitating transcriptional elongation through guanine-rich coding regions. Hum Mol Genet 24, 1824–1835 (2015).

40. Morozov, V.M. et al. HIRA-mediated loading of histone variant H3.3 controls androgen-induced transcription by regulation of AR/BRD4 complex assembly at enhancers. Nucleic Acids Res 51, 10194–10217 (2023).

41. Tafessu, A. et al. H3.3 contributes to chromatin accessibility and transcription factor binding at promoter-proximal regulatory elements in embryonic stem cells. Genome Biol 24, 25 (2023).

42. Rowland, M.E. et al. Systemic and intrinsic functions of ATRX in glial cell fate and CNS myelination in male mice. Nat Commun 14, 7090 (2023).

43. Tillotson, R. et al. A new mouse model of ATR-X syndrome carrying a common patient mutation exhibits neurological and morphological defects. Hum Mol Genet 32, 2485–2501 (2023).

44. Floyd, W. et al. Atrx deletion impairs CGAS/STING signaling and increases sarcoma response to radiation and oncolytic herpesvirus. J Clin Invest 133 (2023).

45. Lorenzi, F. et al. ATRX mutations mediate an immunogenic phenotype and macrophage infiltration in neuroblastoma. Cancer Lett 613, 217495 (2025).

46. Wang, X., Zhao, Y., Zhang, J. & Chen, Y. Structural basis for DAXX interaction with ATRX. Protein Cell 8, 767–771 (2017).

47. Li, M. et al. Dynamic regulation of transcription factors by nucleosome remodeling. Elife 4 (2015).

48. Kadoch, C. et al. Dynamics of BAF-Polycomb complex opposition on heterochromatin in normal and oncogenic states. Nat Genet 49, 213–222 (2017).

49. Goldberg, A.D. et al. Distinct factors control histone variant H3.3 localization at specific genomic regions. Cell 140, 678–691 (2010).

50. Wirbelauer, C., Bell, O. & Schubeler, D. Variant histone H3.3 is deposited at sites of nucleosomal displacement throughout transcribed genes while active histone modifications show a promoter-proximal bias. Genes Dev 19, 1761–1766 (2005).

51. Volkman, H.E., Cambier, S., Gray, E.E. & Stetson, D.B. Tight nuclear tethering of cGAS is essential for preventing autoreactivity. Elife 8 (2019).

52. Kujirai, T. et al. Structural basis for the inhibition of cGAS by nucleosomes. Science 370, 455–458 (2020).

53. Uggenti, C. et al. cGAS-mediated induction of type I interferon due to inborn errors of histone pre-mRNA processing. Nat Genet 52, 1364–1372 (2020).

54. Guey, B. et al. BAF restricts cGAS on nuclear DNA to prevent innate immune activation. Science 369, 823–828 (2020).

55. Rice, G.I. et al. Assessment of interferon-related biomarkers in Aicardi-Goutieres syndrome associated with mutations in TREX1, RNASEH2A, RNASEH2B, RNASEH2C, SAMHD1, and ADAR: a case-control study. Lancet Neurol 12, 1159–1169 (2013).

56. Schindelin, J., et al. Fiji: an open-source platform for biological-image analysis. Nat Methods 9, 676–682 (2012).

57. Subramanian, A. et al. Gene set enrichment analysis: a knowledge-based approach for interpreting genome-wide expression profiles. Proc Natl Acad Sci U S A 102, 15545–15550 (2005).

58. Liberzon, A. et al. The Molecular Signatures Database (MSigDB) hallmark gene set collection. Cell Syst 1, 417–425 (2015).

59. Chen, E.Y. et al. Enrichr: interactive and collaborative HTML5 gene list enrichment analysis tool. BMC Bioinformatics 14, 128 (2013).

60. Cock, P.J. et al. Biopython: freely available Python tools for computational molecular biology and bioinformatics. Bioinformatics 25, 1422–1423 (2009).

61. Chen, S., Zhou, Y., Chen, Y. & Gu, J. fastp: an ultra-fast all-in-one FASTQ preprocessor. Bioinformatics 34, i884–i890 (2018).

62. Dobin, A. et al. STAR: ultrafast universal RNA-seq aligner. Bioinformatics 29, 15–21 (2013).

63. Patro, R., Duggal, G., Love, M.I., Irizarry, R.A. & Kingsford, C. Salmon provides fast and bias-aware quantification of transcript expression. Nat Methods 14, 417–419 (2017).

64. Love, M.I., Huber, W. & Anders, S. Moderated estimation of fold change and dispersion for RNA-seq data with DESeq2. Genome Biol 15, 550 (2014).

65. Ewels, P.A. et al. The nf-core framework for community-curated bioinformatics pipelines. Nat Biotechnol 38, 276–278 (2020).

66. Stephens, M. False discovery rates: a new deal. Biostatistics 18, 275–294 (2017).

67. Buenrostro, J.D. et al. Single-cell chromatin accessibility reveals principles of regulatory variation. Nature 523, 486–490 (2015).

68. Afgan, E. et al. The Galaxy platform for accessible, reproducible and collaborative biomedical analyses: 2018 update. Nucleic Acids Res 46, W537–W544 (2018).

