## Supplemental figures for "ATRX Deficiency Drives Aberrant Type I Interferon Signalling Through cGAS-Dependent Transcriptional Dysregulation"

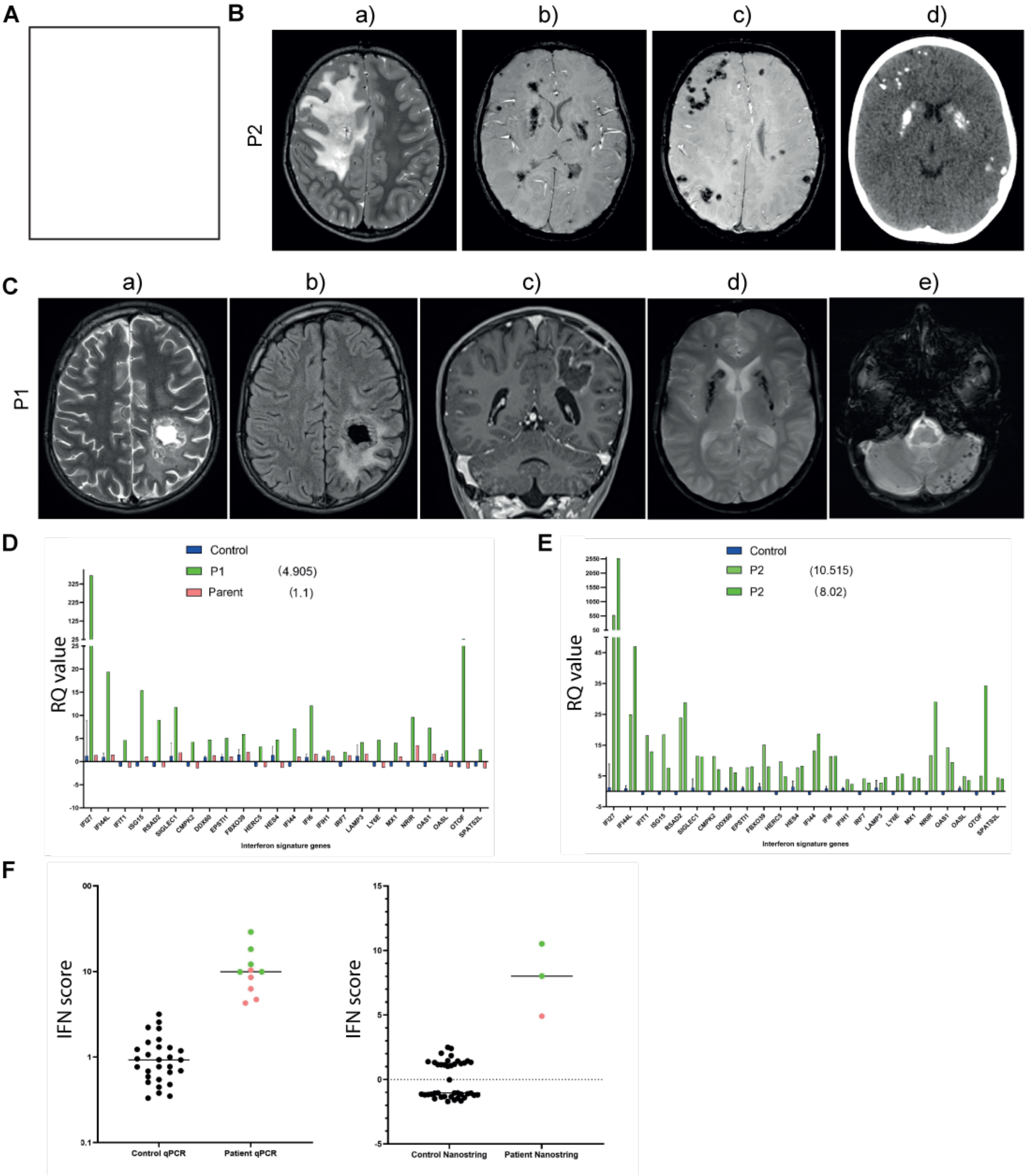


**figure S1. Clinical features and interferon scores in P1 and P2**

(A) Blanck picture. (B) Cerebral imaging in P2: MRI imaging shows a large area of abnormal white matter signal in the right fronto-parietal region, with a cystic area within it (a) (axial T2), and extensive, bilateral but asymmetric, mineralisation in the basal ganglia (b, c) (axial SWI), which was confirmed to be calcification by CT (d). (C) Cerebral imaging in P1: MRI imaging demonstrates a large cystic lesion in the left parietal region with heterogeneous signal intensity (a: axial T2, b: axial FLAIR) with surrounding contrast enhancement (c: coronal post-contrast T1). There is dense mineralisation of the basal ganglia bilaterally (d: axial GRE) and in the left cerebellar hemisphere (e: axial GRE). (D, E) Relative quantification (RQ) values of 24 interferon stimulated genes (ISGs) measured on a NanoString platform in whole blood of P1 (D) and P2 (E). Number in bracket is the interferon (IFN) score, with scores above 2.75 designated as positive. P2 was tested on 2 occasions. Dark blue bars represent the composite data of 27 controls. Note that the IFN score in whole blood of a parent of P1 was normal on the single occasion tested. (F) Left, dot plot showing the IFN score calculated from the median fold change of six ISGs (*IFI27, IFI44L, IFIT1, ISG15, RSAD2, SIGLEC1*) measured by qPCR in whole blood compared to the median of 29 healthy controls, with an abnormal IFN score being defined as greater than +2 s.d. above the mean of the control group, which is set at 2.4. Right, dot plot representation of the data shown in Figure S1D-E. Red = P1, green = P2.

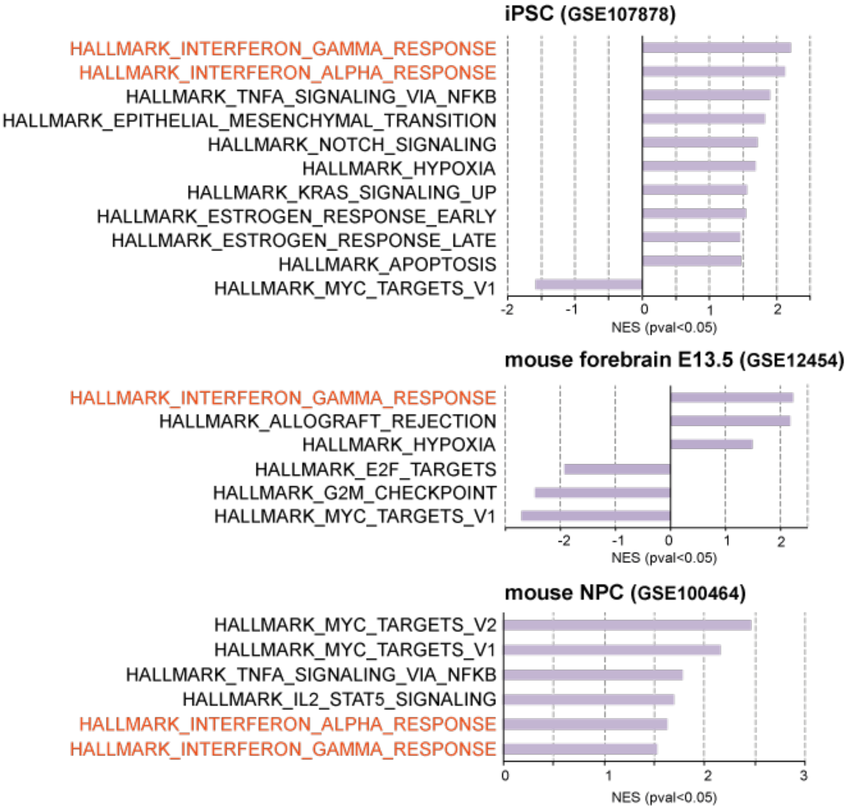


**figure S2. Gene set enrichment analysis (GSEA) of publicly available transcriptomic datasets relating to three ATRX deficient models**

Normalised enrichment scores (NES) of GSEA values with a p value < 0.05 of RNA-Seq data in *ATRX*-null induced pluripotent stem cells (iPSCs) (top), microarray data in *Atrx*-null mouse forebrain (middle) and RNA-Seq data in *Atrx*-null mouse neuronal progenitor cells (NPCs) (bottom). Interferon-related pathways are highlighted in orange. Note: genes from the interferon gamma dataset overlap with the interferon alpha dataset. Transcriptional data were obtained from the Gene Expression Omnibus (GEO) repository, with the accession numbers GSE107878[1], GSE12454[2] and GSE100464[3] as indicated above each data set.

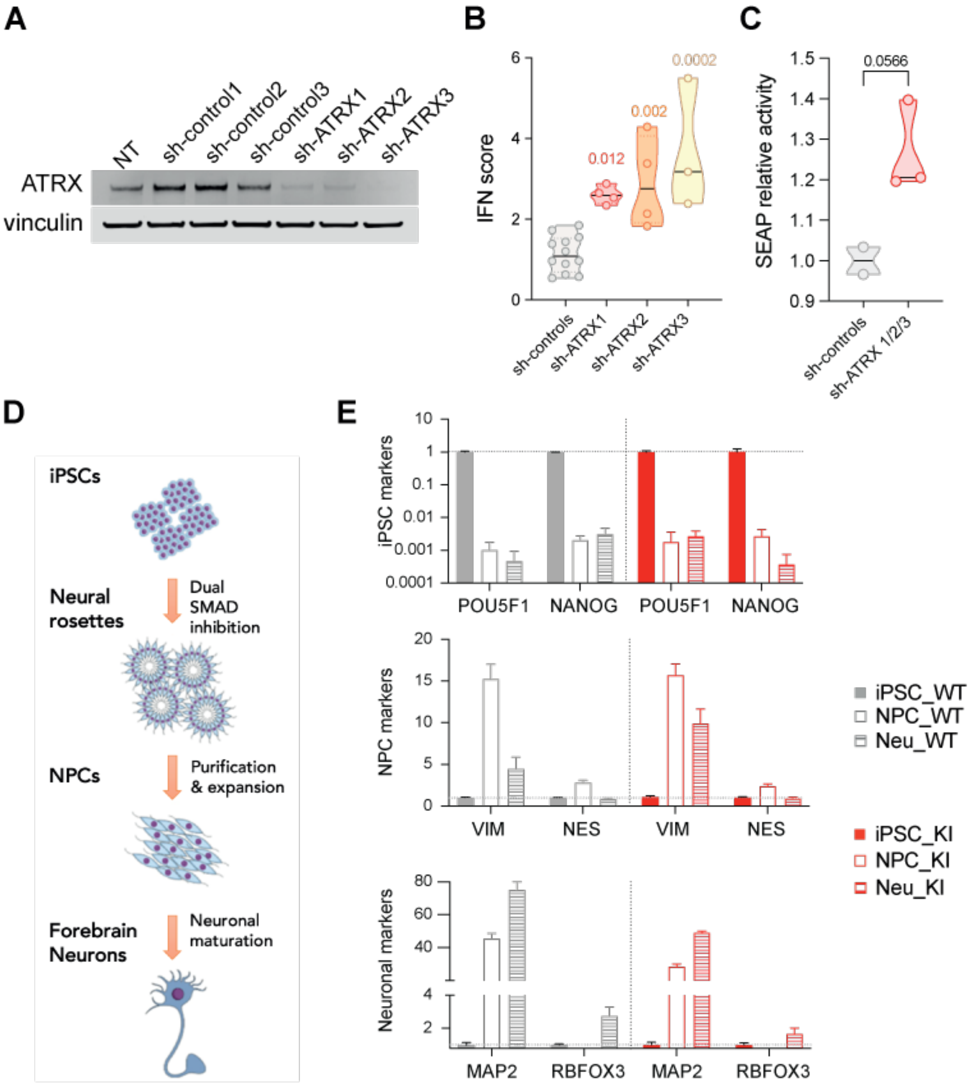


**figure S3. Characterisation of ATRX deficient cell lines and neural differentiation markers**

(A) ATRX protein expression assessed by western blot in THP-1 cells following knock-down of *ATRX* using three short-hairpin (sh) RNAs. (B) Interferon (IFN) score (median expression of *IFI27, RSAD2, ISG15, IFI44L, IFIT1*) in THP-1 cells five days post-transduction with control short-hairpin (sh) or three different shRNAs targeting *ATRX* (sh-ATRX), [three to four independent experiments, black bars represent the median, ordinary one-way ANOVA, Dunnett's multiple comparisons test, individual p values are indicated]. (C) Quantification of secreted type I interferon activity in supernatant from THP-1 cells upon knock-down of *ATRX* by sh-RNA (day 8 post-transduction), measured by secreted embryonic alkaline phosphatase (SEAP) assay in HEK-Blue$^{TM}$ ISG reporter cells [one experiment using three different shRNAs targeting *ATRX*, unpaired t-test]. (D) Schematic of differentiation of human iPSCs to NPCs and then to forebrain neurons following neural induction and neuronal maturation using an established StemCell protocol. (E) Relative gene expression levels of stem cell and neural subtype markers measured by 3' end mRNA sequencing in iPSCs, NPCs and neurons (Neu) WT or *ATRX* KI for the Y1758C substitution.

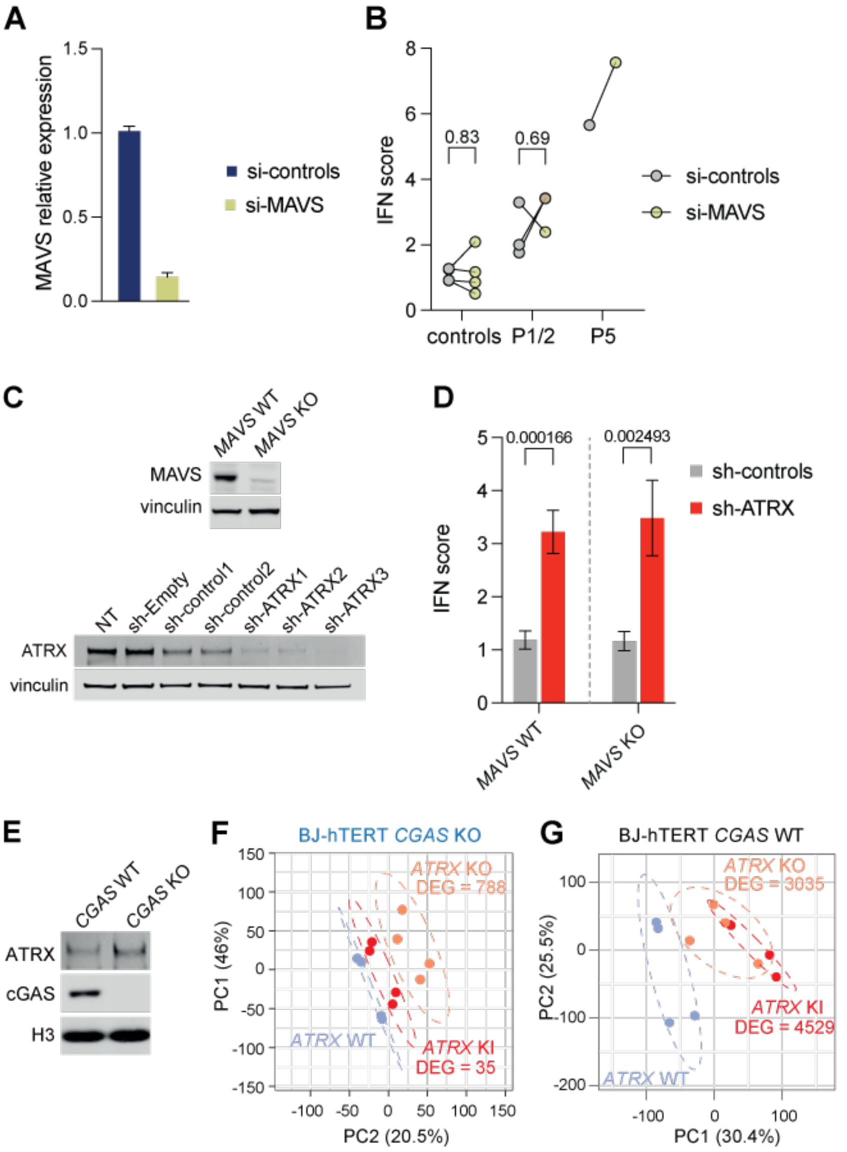


**figure S4. Interferon signalling pathway analysis**

(A) Efficiency of *MAVS* RNA knock-down in primary fibroblasts. (B) Effect on the interferon (IFN) score (median expression *TNFAIP3, RSAD2, Mx1, ISG15, IFI44L, IFIT1, IFI6*) of si-RNA knock-down of *MAVS* in primary fibroblasts from two controls, Y1758C ATRX mutant (P1 and P2) and 'classical' ATRX (P5) mutant fibroblasts, [two independent experiments, dots represent individual repeats, ratio paired t tests; q values are indicated]. (C) Protein expression assessed by western blot of MAVS (top) and ATRX (bottom) in THP-1 cells KO for *MAVS*, following knock-down of *ATRX* using three short-hairpin (sh) RNAs. (D) Interferon (IFN) score (median expression of *IFI27*, *IFIT1*, *RSAD2*, *ISG15*) in THP-1 cells five days post-transduction with control shRNAs (sh-controls) or three different shRNAs targeting *ATRX* (sh-ATRX) [four independent experiments, mean +/- SEM, Mann-Whitney tests, q values are indicated]. (E) Protein levels assessed by western blot in nuclear fractions of BJ-hTERT clones WT or KO for *CGAS*. (F-G) Principal component (PC) analysis of RNA-Seq data derived from four BJ-hTERT fibroblasts clones KO (F) or WT (G) for *CGAS*, and WT, KI or KO for *ATRX*. The number of differentially expressed genes (DEG, p adjusted < 0.1) identified between WT and KI or WT and KO *ATRX* cells are noted on the graph. All annotated human genes were analysed.

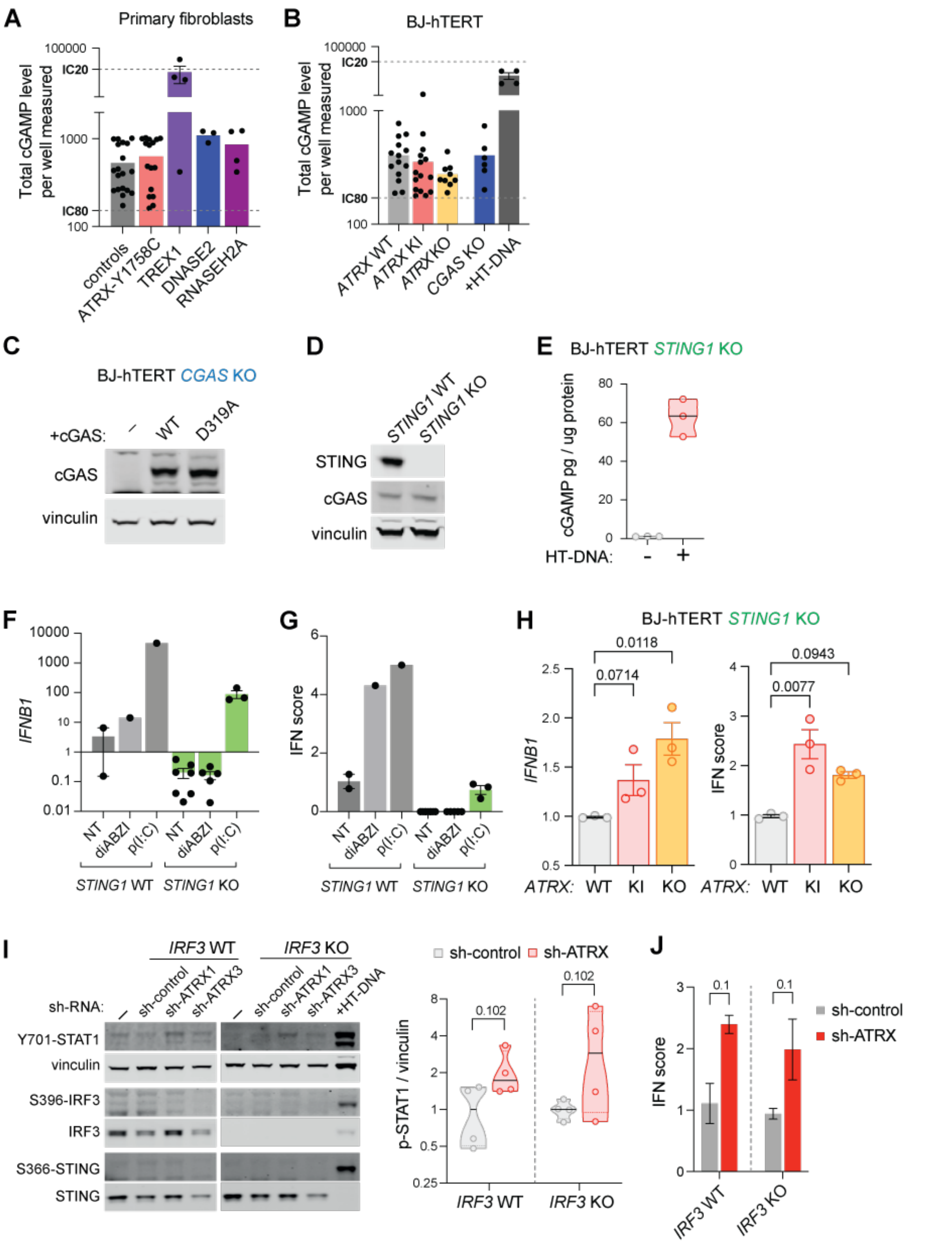


**figure S5. Canonical DNA sensing pathway analysis**

(A-B) Graphs illustrating total cGAMP levels measured by ELISA in wells containing total cellular lysate obtained from primary fibroblasts (A) or BJ-hTERT clones (B): dotted lines indicate the level of IC20 and IC80, representing the upper and lower limits of detection of the standard curve, respectively. Dots represent individual measurements and relate to Figure 4A and 4B. (C) Western blot showing protein levels of transfected WT or catalytic inactive (D319A mutant) cGAS constructs. (D) Protein levels assessed by western blot of BJ-hTERT fibroblasts clones WT or KO for *STING1*. (E) cGAMP levels measured in BJ-hTERT fibroblasts KO for *STING1* unstimulated (-) or stimulated (+) with HT-DNA for four hours, [one experiment, three replicates]. (F-G) Interferon B1 (*IFNB1*) gene expression (F) and interferon (IFN) score (median expression of *IFI27, IFI6, IFIT1, OAS1, ALDH1A1, IFIH1*) (G) in BJ-hTERT fibroblast WT or KO for *STING1*, unstimulated (NT), and stimulated with STING agonist diABZI or the MAVS agonist p (I:C) (synthetic analogue of double-stranded RNA polyinosinic-polycytidylic acid). (H) IFN score (median expression of *IFI27, IFI6, IFIT1, OAS1, ALDH1A1, IFIH1*) (left) and relative expression of *IFNB1* (right) measured by qPCR in BJ-hTERT fibroblasts KO for *STING1*, and WT, KI or KO for *ATRX* [three independent clones repeated three times, average +/- SEM, Kruskal-Wallis test, two-stage linear step-up procedure of Benjamin, Krieger and Yekutieli]. (I) Left: Immunoblot of phosphorylated STAT1 (Y701), STING (S366), phosphorylated IRF3 (S396), total IRF3 and STING in THP-1 cells WT or KO for *IRF3*, transduced with control sh-RNAs or sh-RNAs targeting *ATRX*. The last lane relates to WT THP-1 cells stimulated with HT-DNA for four hours; Right: Quantification of STAT1 phosphorylation level over vinculin, [two independent experiments, circles are individual repeats, black bars indicate the means, Kruskal-Wallis test, two-stage linear step-up procedure of Benjamin, Krieger and Yekutieli, q values are indicated] (J) IFN score (median expression of *IFI27*, *IFIT1*, *RSAD2*, *ISG15*) in THP-1 cells five days post-transduction with control sh-RNA (sh-control) or two different sh-RNAs targeting *ATRX* (sh-ATRX) [one independent experiment, mean +/- SEM, Mann-Whitney tests, q values are indicated].

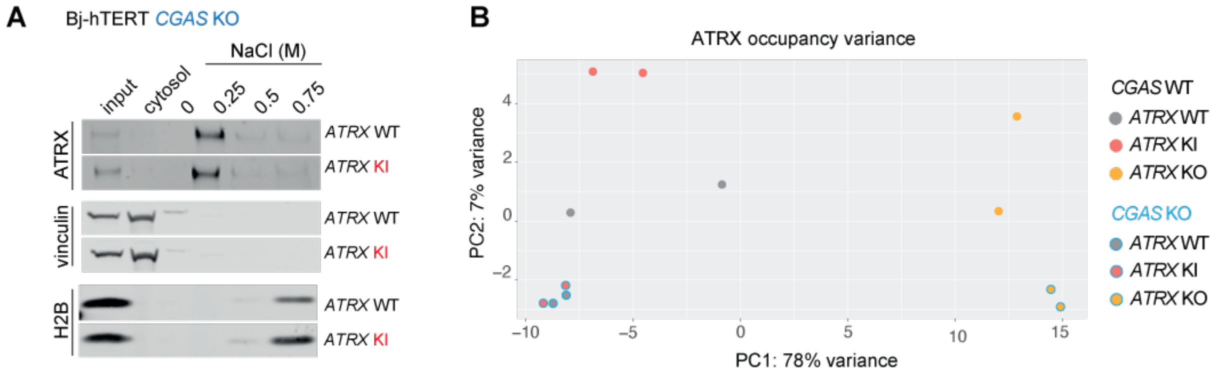


**figure S6. Characterization of ATRX Y1758C chromatin binding in cGAS null cells**

(A) Subcellular fractionation of BJ-hTERT fibroblasts WT or KI for *ATRX*. Cells were separated into cytosolic and nuclear fractions, followed by sequential elution of nuclear pellets with the indicated concentrations of NaCl. ATRX, vinculin and H2B were assessed by western blot. (B) Principal component (PC) analysis comparing the ATRX genomic binding profiles between indicated genotypes.

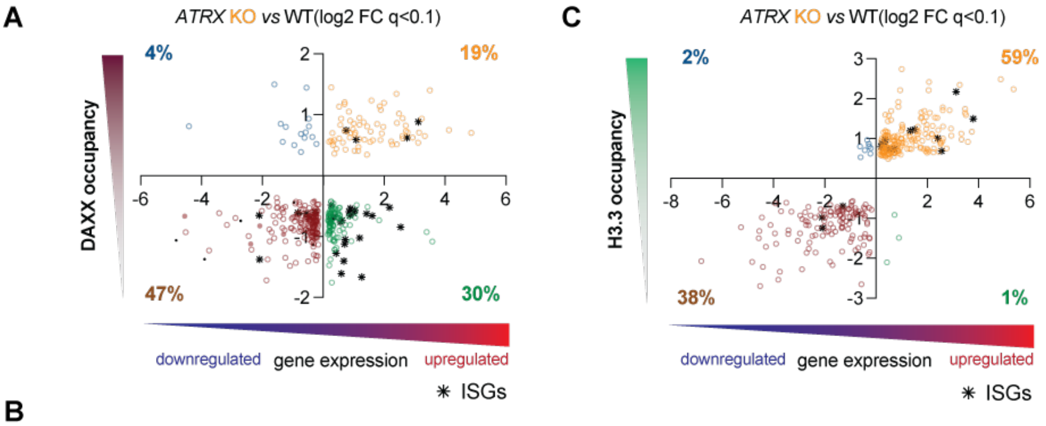


B

Pathway enrichment analysis of genes showing decreased levels of DAXX and upregulated expression in ATRX deficient cells

| | Term | P value | Adjusted P value | Combined Score |
|---|---|---|---|---|
| ATRX KI *vs* WT | Interferon Alpha Response | 1.3E-18 | 4.9E-17 | 1244 |
| | Interferon Gamma Response | 9.8E-17 | 1.8E-15 | 592 |
| ATRX KO *vs* WT | Interferon Alpha Response | 4.2E-13 | 1.4E-11 | 758 |
| | Interferon Gamma Response | 2.2E-09 | 3.5E-08 | 239 |

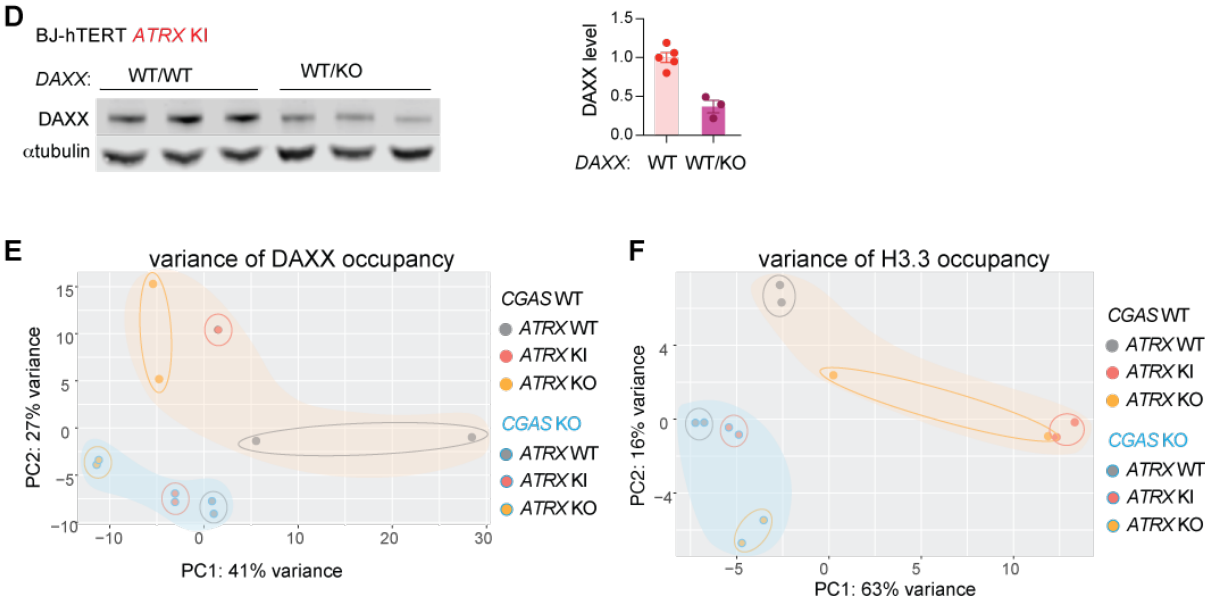


**figure S7. Loss of ATRX function disrupts DAXX and H3.3 occupancy at genes**

(A) Dot plot of genes exhibiting significant (adjusted p value (q) < 0.1, x-axis) changes in both transcription and in DAXX occupancy, in *ATRX* KO as compared to WT cells. Quadrant colour coding is as follows: blue = genes enriched in DAXX and downregulated in expression; orange = genes enriched in DAXX and upregulated in expression; brown = genes depleted of DAXX and downregulated in expression; green = genes depleted of DAXX and upregulated in expression; ISGs are denoted separately by a black asterisk, while other genes are represented by empty circles. (B) Table showing the top two significant terms of the pathway enrichment analysis of genes whose expression is increased in cells KI or KO for *ATRX* and exhibiting decreased levels of DAXX. (C) Dot plot of genes exhibiting significant (adjusted p value (q) < 0.1, x-axis) changes in both transcription and in H3.3 occupancy, in *ATRX* KO as compared to WT cells, with the colour coding as described in (A). (D) Western blot (left) and total protein quantification (right) of DAXX in BJ-hTERT fibroblasts KI for *ATRX* and WT or heterozygous KO (WT/KO) for *DAXX*, [circles represent individual clones]. (E-F) Principal component (PC) analysis comparing DAXX (E) and H3.3 (F) genomic binding profiles between indicated genotypes.

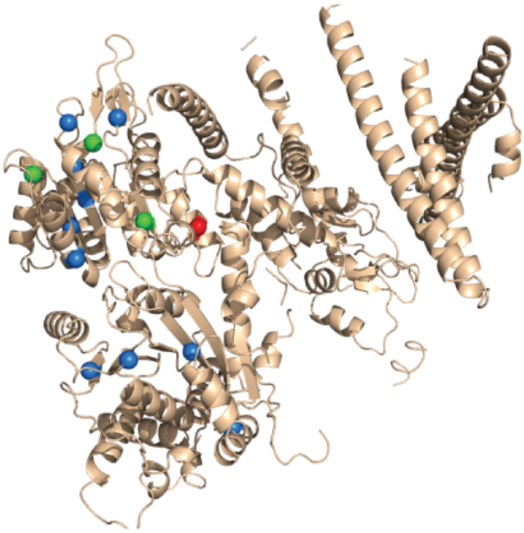

**figure S8. Structural mapping of ATRX mutations**

Mapping of the Y1758C ATRX mutation (in red), three mutations (V1552F, K1650N and L1746S: in green) described by Mitson et al. [4] to affect ATRX protein function, and the sites of other mutations that strongly reduced protein levels described in the same study (blue). L1746S lies in a conserved helicase motif, and Y1758C at the start of helix α12 (using numbering from the homologous Rad54 structure). Mutations were mapped onto a model of the ATRX structure from the AlphaFold Protein Structure Database (alphafold.ebi.ac.uk), based upon UniProtID P46100. The structure was visualised using PyMol, excluding low-confidence/disordered regions (pLDDT ≤ 50).

**Supplementary references**


1. Deneault, E. *et al.* Complete Disruption of Autism-Susceptibility Genes by Gene Editing Predominantly Reduces Functional Connectivity of Isogenic Human Neurons. *Stem Cell Reports* **11**, 1211-1225 (2018).

2. Levy, M.A., Fernandes, A.D., Tremblay, D.C., Seah, C. & Berube, N.G. The SWI/SNF protein ATRX co-regulates pseudoautosomal genes that have translocated to autosomes in the mouse genome. *BMC Genomics* **9**, 468 (2008).

3. Danussi, C. *et al.* Atrx inactivation drives disease-defining phenotypes in glioma cells of origin through global epigenomic remodeling. *Nat Commun* **9**, 1057 (2018).

4. Mitson, M., Kelley, L.A., Sternberg, M.J., Higgs, D.R. & Gibbons, R.J. Functional significance of mutations in the Snf2 domain of ATRX. *Hum Mol Genet* **20**, 2603-2610 (2011).
